# ApoE isoforms establish distinct antioxidant trajectories across the aging brain

**DOI:** 10.64898/2026.11.06.765000

**Authors:** Sarah J. Shemtov, Mafalda Cacciottolo, Amy Christensen, Boyang Li, Amin Haghani, Michelle C. Rice, Herbert Anson, Claire S. Chung, Whitaker Cohn, Hongqiao Zhang, Wendy J. Mack, Annie Hiniker, Changhan David Lee, Todd E. Morgan, Elizabeth Head, Henry Jay Forman, Ryo Higuchi-Sanabria, Christian J. Pike, Hussein Yassine, Marc Vermulst, Caleb E. Finch, Max A. Thorwald

## Abstract

*APOE ε4* is the strongest common genetic risk factor for late-onset Alzheimer’s disease, but how ApoE isoforms shape physiology before disease remains unclear. Across the ancestral-to-derived ApoE4→ApoE3→ApoE2 sequence, we examined antioxidant regulation during aging. Young humanized ApoE4 mice had lower antioxidant protein abundance and oxidized phospholipid reduction before detectable oxidative damage, yet accumulated lipid peroxidation products with age despite antioxidant remodeling. ApoE4 cells mounted the strongest antioxidant transcriptional response to sulforaphane but accumulated less protein from targets supporting proteostasis, glutathione synthesis and ferroptosis suppression, alongside altered eIF2α signaling and translation kinetics. Antioxidant transcription shifted from an ApoE4 bias in youth toward ApoE2 with age, and adult APOE4-to-APOE2 switching redirected expression in astrocytes and microglia. Cognitively normal human brains showed regional reductions in nuclear Nrf2 and antioxidant proteins, with transcriptomic analyses supporting context-dependent organization. ApoE isoforms differentially couple stress-responsive transcription to protein output, linking age-dependent antioxidant adaptation to neurodegenerative vulnerability.

## Introduction

*APOE* is the strongest genetic modifier of late-onset Alzheimer’s disease (AD): ε4 increases risk and advances onset in a dose-dependent manner, whereas ε2 lowers risk and is associated with longevity^1,2^. Yet the conventional description of *APOE* ε4 as deficient, *APOE* ε3 as neutral and *APOE* ε2 as protective is a disease-centered hierarchy rather than a biological explanation. At residues 112 and 158, ApoE4 retains the ancestral arginine, whereas ApoE3 and ApoE2 carry successive cysteine substitutions^3^. Chimpanzee ApoE is correspondingly closer in sequence and aspects of function to human ApoE4, although additional species-specific substitutions prevent their equivalence^4^. The ApoE4→ApoE3→ApoE2 sequence therefore provides an evolution-informed framework for testing how the human isoforms reorganize cellular physiology without characterizing ApoE4 as a deficient form of ApoE3.

The persistence of *APOE* ε4 in some populations despite its late-life disease associations indicates that its effects depend on biological and environmental context. Among the Tsimane, Bolivian forager-horticulturalists with high pathogen exposure, *APOE* ε4 is associated with better cognition under high parasitic load and greater fecundity^5,6^. *APOE ε4* is also associated with reduced melanoma progression and greater antitumor immunity, opposing the APOE2-to-APOE4 relationship observed in AD^7^. Conversely, *APOE ε2* is associated with longevity independently of AD neuropathology, and human proteomic analyses link APOE2 to cellular maintenance and anti-inflammatory signatures before clinical disease^2,8^. These observations support a model in which the isoforms differentially weigh inflammatory defense and long-term cellular maintenance, with rebalancing as metabolic capacity declines and sterile inflammation rises with age or environmental context.

The aging brain provides a stringent setting to test this model. High metabolic demand, oxidation-prone membrane lipids and accumulation of redox-active iron favor lipid peroxidation and ferroptotic injury^9,10^. Protection requires coordinated glutathione synthesis, phospholipid-hydroperoxide detoxification and other parallel ferroptosis-suppressive defenses. We previously found iron-associated lipid peroxidation, lower ferroptosis-suppressive proteins and their corresponding activities in human AD brain, with prominent deficits among *APOE ε4* carriers^11^. Complementing this evidence, ApoE2 and ApoE3 Christchurch promote ABCA7-dependent clearance of oxidized neuronal lipids, whereas ApoE4 increases lipid-peroxide burden and ferroptotic vulnerability^12^. ApoE-dependent stress adaptation also extends beyond lipid detoxification: ApoE2 promotes DNA repair and resistance to neuronal senescence, while young ApoE4 mice exhibit eIF2α phosphorylation otherwise associated with aging^13,14^. Chronic low-grade inflammation disproportionately increases AD risk among APOE ε4 carriers^15^, while Nrf2 activates antioxidant defense and suppresses inflammatory transcription^16,17^. Comparing antioxidant, inflammatory and pro-oxidant expression can therefore distinguish selective antioxidant remodeling from broader changes in stress signaling.

Evidence linking ApoE genotype to Nrf2 and antioxidant balance remains limited. ApoE4 -targeted-replacement mice have lower hepatic nuclear Nrf2 and reduced Nrf2-target expression than ApoE3 mice^18^, while brain transcriptomics show that age interacts strongly with ApoE genotype and sex^19^. APOE effects are also cell-type dependent: human single-nucleus analyses identify distinct isoform relationships across neural and vascular populations^20^, and isogenic human models demonstrate cell-autonomous and non-cell-autonomous dysregulation between APOE4 astrocytes and microglia^21^. Most importantly, inducible APOE4-to-APOE2 replacement demonstrates that ApoE-dependent brain states remain modifiable in adulthood and that astrocyte-restricted switching can alter multiple CNS cell types^22^. How aging and adult allelic switching reshape ApoE-dependent antioxidant regulation remains unknown. Induced RNAs must be processed and translated to produce protective proteins, yet ApoE’s contribution to this coupling has not been tested directly.

Here we investigated ApoE-dependent antioxidant states before overt neuropathology. We integrated two humanized mouse models and primary glia with aging and adult allelic-switch transcriptomics, SFN stimulation, and human regional and single-nucleus analyses. These experiments examined the relationship between antioxidant transcription, protein accumulation and functional defense across age and cellular context and tested the capacity of adult isoform replacement to redirect antioxidant expression.

## Results

### ApoE isoforms establish region- and age-dependent antioxidant states

Membrane phospholipids are among the most oxidation-prone molecules in the brain, and only two enzymes, GPx4 and Prdx6, reduce the resulting phospholipid hydroperoxides directly. Both require glutathione, synthesized by the rate-limiting GCLC-GCLM complex. We therefore measured all four proteins in two independent humanized ApoE targeted-replacement models: CAF-TR mice at 6 and 22 months and ApoE-TR mice at 6 months (**Fig. 1A**).

**Figure 1.**
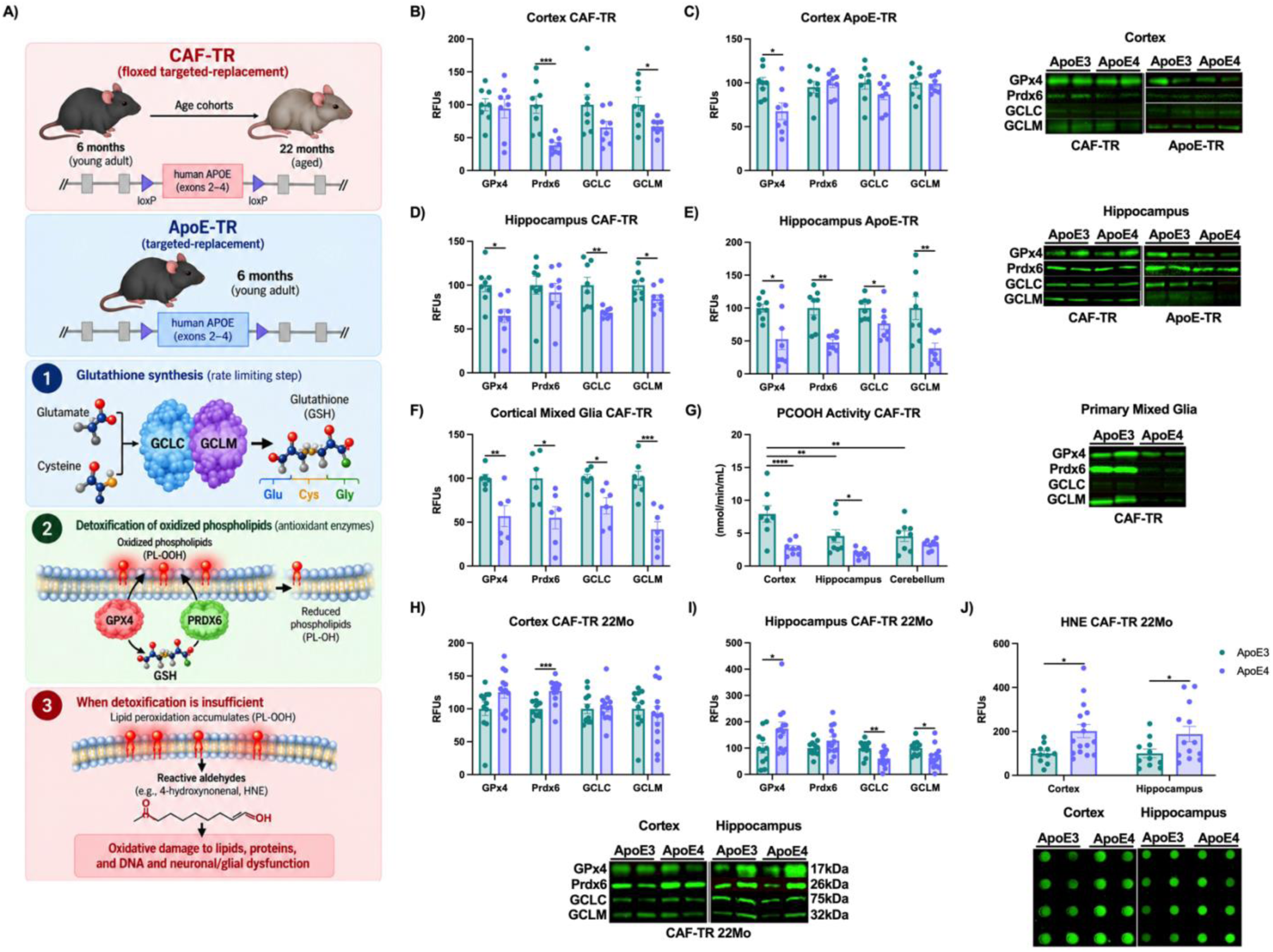
ApoE genotype alters the glutathione-dependent phospholipid antioxidant system across brain aging. **A)** Schematic of the two humanized ApoE targeted-replacement mouse models and antioxidants critical to ferroptosis examined in this study. CAF-TR and ApoE-TR mice independently replace the endogenous mouse Apoe coding sequence with human APOE; the CAF-TR allele is additionally flanked by loxP sites and was examined at 6 and 22 months of age, whereas ApoE-TR mice were examined at 6 months. GCLC and GCLM form glutamate-cysteine ligase (GCL), the rate-limiting enzyme in glutathione (GSH) synthesis. GSH supports GPx4- and Prdx6-mediated reduction of oxidized phospholipids, whereas insufficient detoxification promotes lipid peroxidation and formation of reactive aldehydes, including 4-hydroxynonenal (HNE). Protein abundance (n=4/sex/genotype) of GPx4, Prdx6, GCLC and GCLM in 6-month-old **B)** CAF-TR cortex, **C)** ApoE-TR cortex, **D)** CAF-TR hippocampus and **E)** ApoE-TR hippocampus. Within each mouse model, brain regions were collected from the same animals. **F)** GPx4, Prdx6, GCLC and GCLM protein abundance in primary cortical mixed glia from neonatal CAF-TR mice. **G)** Oxidized phosphatidylcholine (PCOOH) reducing activity in cortex, hippocampus and cerebellum from 6-month-old CAF-TR mice; cortex and hippocampus were from the same animals analyzed in panels B and D. The same proteins in 22-month-old CAF-TR (n=4-8/sex/genotype) **H)** cortex and **I)** hippocampus. **J)** HNE adducts in cortex and hippocampus from the same 22-month-old CAF-TR mice. Representative immunoblots are shown adjacent to or below the corresponding quantification. Protein abundance and HNE are expressed as relative fluorescence units (RFUs). Data are shown as mean±SEM. B-F,H-J) by two-tailed unpaired t-test or Mann-Whitney test as appropriate. G) by two-way ANOVA with ApoE isoform and brain region as factors followed by Tukey’s multiple-comparisons test. *p<0.05, **p<0.01, ***p<0.001, ****p<0.0001.

At 6 months, ApoE4-associated reductions were evident in both models but varied by brain region. In CAF-TR cortex, Prdx6 and GCLM were lower in ApoE4 mice, whereas GPx4 and GCLC were unchanged (**Fig. 1B**). In ApoE-TR cortex, GPx4 was 33% lower in ApoE4 mice, whereas Prdx6, GCLC and GCLM were unchanged (**Fig. 1C**). The hippocampal phenotype was broader, with 35%, 32% and 16% less GPx4, GCLC and GCLM, respectively in ApoE CAF-TRs (**Fig. 1D**). The same regional selectivity was observed in ApoE-TR mice with 23-69% less of all four proteins for the hippocampus (**Fig. 1E**). To determine if these changes were present early on, we cultured primary cortical mixed glia from neonatal CAF-TR mice, which had at least 30% less GPx4, Prdx6, GCLC and GCLM (**Fig. 1F**).

Having established that the proteins themselves were reduced, we next evaluated whether this altered their functional capacity. To do this, we measured the enzymatic reduction of oxidized phosphatidylcholine (PCOOH). PCOOH activity was 66% lower in ApoE4 CAF-TR cortex and 58% lower in hippocampus but was unchanged in cerebellum mirroring the neuropathological burden observed during AD (**Fig. 1G**). Total GPx activity was unchanged in young cortex and hippocampus, distinguishing impaired phospholipid hydroperoxide reduction from a generalized loss of GPx activity, whereas cortical SOD activity was 30% lower in ApoE4 mice (**Ext. Fig. 1A,B**). Because reduced antioxidant capacity could either precede or follow oxidative injury, we measured markers of accumulated damage at the same age. HNE adducts and nitrotyrosine, reflecting lipid peroxidation and protein nitration, did not differ by genotype (**Ext. Fig. 1C,D**), placing the ApoE4 deficit in antioxidant protein and activity upstream of detectable oxidative damage rather than downstream of it. The phenotype was also tissue-selective rather than systemic, as GPx4 was increased in heart, liver and lung of ApoE4 mice while the midbrain showed lower Prdx6 and GCLC (**Ext. Fig. 1E-H**).

We next asked whether this early antioxidant deficit persisted during advanced aging. In 22-month-old CAF-TR cortex, Prdx6 was 27% higher in ApoE4 mice, while the other proteins were unchanged (**Fig. 1H**). In hippocampus, GPx4 was 73% higher, whereas GCLC and GCLM remained 40% and 37% lower, respectively (**Fig. 1I**). Neither PCOOH nor total GPx activity differed by genotype in either region (**Ext. Fig. 1I,J**). Despite this convergence, HNE was 56% higher in ApoE4 cortex and 88% higher in hippocampus (**Fig. 1J)**. The complete pattern across models, tissues and ages is summarized in **Ext. Fig. 1K**. Thus, ApoE4 produced an early deficit in phospholipid antioxidant machinery and function that preceded detectable damage, followed by partial remodeling with age that failed to prevent lipid peroxidation.

Because Nrf2 activates these defenses through nuclear accumulation and binding of the electrophile response element (EpRE), we asked whether its promoter occupancy tracked this age-dependent reduced protein phenotype. Young ApoE4 cortex contained less nuclear Nrf2 in both mouse models, whereas cytosolic Nrf2 was higher in ApoE-TR and lower in CAF-TR mice; ApoE4 primary glia had twofold more cytosolic Nrf2 without a significant nuclear increase (**Ext. Fig. 2A-D**). Nrf2 occupancy was also lower at the Prdx6 promoter in young ApoE4 cortex, with a similar trend at the GPx4 promoter (**Ext. Fig. 2E,F**). By 22 months, nuclear Nrf2 was instead higher in ApoE4 cortex (**Ext. Fig. 2D**). Thus, reduced nuclear localization and promoter engagement accompanied the early ApoE4 deficit in phospholipid hydroperoxide defense, whereas increased nuclear Nrf2 with age did not prevent lipid peroxidation.

### ApoE-dependent antioxidant transcription reverses across brain aging

The age-dependent protein remodeling raised the question of whether ApoE isoforms establish fixed antioxidant states or whether their transcriptional relationship changes across lifespan. We addressed this using published cortical RNA-sequencing data from 143 ApoE-TR mice at 3, 12 and 24 months^19^. To examine expression along the ancestral-to-derived ApoE4→ApoE3→ApoE2 trajectory, genotype was modeled ordinally, assigning them −1, 0 and +1 respectively. A single coefficient per gene then captures which end of that axis the gene favors, with negative values indicating higher expression toward ApoE4 and positive values toward ApoE2 (**Fig. 2A**). We applied this to antioxidant, inflammatory and pro-oxidant predefined gene sets so that changes in antioxidant regulation could be distinguished from a general shift in stress signaling.

**Figure 2.**
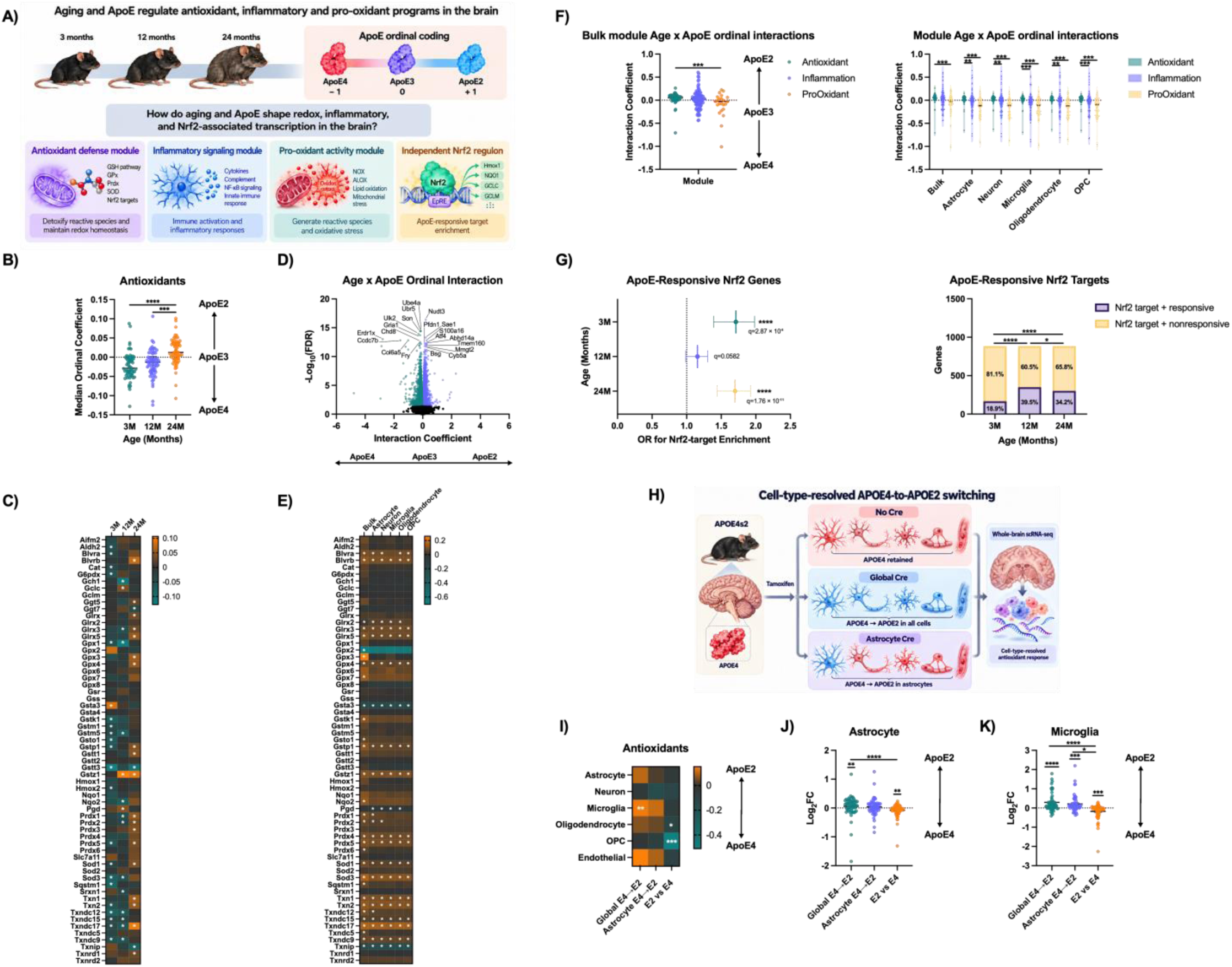
ApoE-dependent antioxidant transcription reverses across aging and is reconfigured by adult APOE4-to-APOE2 switching. Cortical RNA-sequencing data from 143 ApoE-TF mice at 3, 12 and 24 months (n=15-16/genotype/sex) were analyzed using ordinal coding: ApoE4=−1, ApoE3=0 and ApoE2=+1. Positive age-specific coefficients indicate ApoE2-biased expression; positive Age x ApoE interaction coefficients indicate a shift toward ApoE2 between 3 and 24 months. **A)** Experimental and analytical framework. **B)** Age-specific ordinal-coefficient distributions for the predefined 62-gene antioxidant panel. **C)** Heatmap of age-specific coefficients for individual antioxidant genes. **D)** Transcriptome-wide 3-to-24-month Age x ApoE ordinal interaction. **E)** Heatmap of antioxidant-gene interaction coefficients in bulk cortex and MuSiC-adjusted cell-type-enriched models. **F)** Antioxidant, inflammatory and pro-oxidant interaction-coefficient distributions in bulk cortex and the indicated cell-type-enriched models. **G)** Enrichment of an independent Nrf2 regulon among ApoE-responsive genes, shown as odds ratios with 95% confidence intervals, and the responsive and nonresponsive fractions of the 884 represented Nrf2 targets at each age. ApoE responsiveness was defined by FDR<0.05 in either categorical ApoE2-versus-ApoE3 or ApoE4-versus-ApoE3 comparison. **H)** Experimental design for adult global or astrocyte-restricted APOE4-to-APOE2 switching in APOE4s2 mice and comparison with constitutive ApoE2-TR and ApoE4-TR mice. **I)** Median antioxidant-gene log_2_ fold changes across cell populations for global-switch versus no-Cre, astrocyte-switch versus no-Cre and ApoE2-TR versus ApoE4-TR comparisons. Positive values indicate higher expression in the first group named. Corresponding gene-level distributions in **J)** astrocytes and **K)** microglia. **L)** APOE ε3/ε3-versus-ε4/ε4 log_2_ fold changes for 50 antioxidant genes in isogenic human iPSC-derived neuron-astrocyte co-cultures, with four independently cultured samples per genotype. Distribution plots show individual genes with median and interquartile range. B) Friedman test with Dunn’s multiple-comparisons test. Gene-level coefficients in C-E) were evaluated using DESeq2 Wald tests with Benjamini-Hochberg correction; heatmap asterisks denote FDR<0.05. F) Two-sided Wilcoxon signed-rank tests against zero and permutation comparisons between panels with Benjamini-Hochberg correction. G) Nrf2-regulon enrichment by two-sided Fisher’s exact tests with Benjamini-Hochberg correction. Age-dependent differences in the responsive fraction of the same 884 Nrf2 targets were assessed by Cochran’s Q test, followed by pairwise McNemar tests with Holm correction. I) CAMERA with Benjamini-Hochberg correction using mouse-level pseudobulk profiles. J,K) Friedman tests with Dunn’s multiple-comparisons tests across contrasts and two-sided Wilcoxon signed-rank tests against zero with Benjamini–Hochberg correction. L) Differential expression by DESeq2; panel distribution by two-sided Wilcoxon signed-rank test with Benjamini-Hochberg correction across predefined panels. *P<0.05, **P<0.01, ***P<0.001, ****P<0.0001.

Within the predefined 62-gene antioxidant panel, expression was biased toward ApoE4 at 3 and 12 months and reversed toward ApoE2 at 24 months (**Fig. 2B,C**). The 24-month distribution differed from both younger ages, which did not differ from each other. Age-specific inflammatory and pro-oxidant patterns, transcription-factor activity and representative genes are shown in **Ext. Fig. 3A-I**. Testing this transition directly as an Age x ApoE genotype interaction, we found that 54 of 62 antioxidant genes shifted toward ApoE2 between 3 and 24 months (Wilcoxon signed-rank *P*=2.6x10^−7^; **Fig. 2D**). Transcriptome-wide enrichment GSEA analyses independently identified reactive oxygen species pathway signaling as ApoE2-biased at 24 months only, reproducing the antioxidant trajectory without relying on the predefined panel (**Ext. Fig. 3J-L**).

To determine whether the bulk-cortex transition extended across major neural cell types, we used MuSiC deconvolution to estimate cell-type-enriched expression profiles. The positive Age x ApoE interaction was reproduced in neuron-, astrocyte-, oligodendrocyte-, microglia- and oligodendrocyte precursor cell (OPC)- enriched profiles, with 42-46 genes per cell type shifting toward ApoE2 (*P*≤1.2x10^−3^; **Fig. 2E, Ext. Fig. 4**), indicating the reversal is not driven by shifting proportions or by one dominant cell type. We next tested whether aging shifted stress-associated transcription broadly or distinguished antioxidant defense from inflammatory and pro-oxidant expression. Bulk median interaction coefficients were +0.056 for antioxidant, +0.028 for inflammatory and −0.028 for pro-oxidant genes. Pro-oxidant coefficients remained negative in every cell-type-enriched model, whereas inflammatory coefficients varied by cell type (**Fig. 2F**). Antioxidant and pro-oxidant distributions differed in bulk cortex and all five cell-type models; antioxidant and inflammatory distributions differed in the cell-type models but not bulk cortex.

Finally, we repeated the analysis with an independent ChIP-seq-derived mouse Nrf2 regulon of 1,055 basal or inducible targets, of which 884 were represented in the ApoE-TR transcriptome. Nrf2 targets were enriched among ApoE-responsive genes at 3 months (167/884; 18.9%; OR=1.71; *P*=2.9x10^−8^) and 24 months (302/884; 34.2%; OR=1.70; *P*=1.8x10^−11^). Despite the larger responsive fraction at 12 months, enrichment relative to the remaining transcriptome did not survive correction (349/884; 39.5%; OR=1.16; *P*=0.06; **Fig. 2G**).

Because ordinal coding assumes an equally spaced relationship, we independently positioned ApoE3 between categorical ApoE4 and ApoE2 expression estimates. Among genes falling between those endpoints, ApoE3 positions centered near the predicted midpoint across gene sets and cell-type-enriched profiles. Genome-wide positional classes redistributed with age, whereas antioxidant and pro-oxidant classes did not; inflammatory classes changed between 3 months and both later ages (**Ext. Fig. 5A-I**).

We next asked whether the antioxidant state remained responsive to ApoE identity in the adult brain. To test this, we reanalyzed whole-brain single-cell RNA sequencing from APOE4s2 mice following global or astrocyte-restricted APOE4-to-APOE2 switching, along with unswitched and constitutive ApoE2-TR and ApoE4-TR controls (**Fig. 2H**). In constitutive ApoE2-TR versus ApoE4-TR mice, antioxidant expression was ApoE4-biased across all six cell populations, reproducing the young ApoE-TR state. In contrast, global switching in adulthood reversed this in astrocytes, microglia, oligodendrocytes and endothelial cells, though not in neurons and OPCs. Astrocyte-restricted switching produced the same pattern, including in microglia, cells whose own genotype did not change (**Fig. 2I-K**), indicating that the ApoE identity of one cell type can shape the antioxidant state of another.

The early ApoE4-biased state also extended to human neural cells. In isogenic iPSC-derived neuron– astrocyte co-cultures, 35 of 50 detected antioxidant genes were more highly expressed in APOE ε4/ε4 than ε3/ε3 cultures, with a median ε3/ε3-versus-ε4/ε4 log_2_ fold change of -0.55 (**Fig. 2L**). The contrasting constitutive and switched states indicate that antioxidant regulation depends on age, cell type and prior isoform exposure.

### ApoE4 produces a strong Nrf2 transcriptional response to sulforaphane

The ApoE4-biased antioxidant transcription in young models contrasted with lower antioxidant protein abundance *in vivo*. We therefore asked how ApoE isoforms shape the response to acute Nrf2 activation. Isoform-expressing U87 cells were treated with sulforaphane (SFN) for 18 hours; cell-model generation, validation and experimental allocation are shown in **Ext. Fig. 6**.

SFN induced canonical Nrf2 targets, including TXNRD1, SLC7A11, SQSTM1, NQO1 and GCLC, in all three ApoE backgrounds (**Fig. 3A-C**). GSEA identified shared reactive oxygen species, xenobiotic metabolism, mTORC1 and p53-associated responses across the three isoforms (**Ext. Fig. 7A-C**). The response was broadest in ApoE4, comprising 5,425 DEGs compared with 5,181 in ApoE3 and 4,234 in ApoE2; gene-state distributions differed across every isoform comparison (**Ext. Fig. 7D**). Within the antioxidant panel, median SFN-induced log_2_ fold changes increased from +0.053 in ApoE2 to +0.070 in ApoE3 and +0.140 in ApoE4 across 55 shared genes (Friedman *P*=0.0077; **Fig. 3D**). The heatmap showed that the stronger ApoE4 response extended across the panel rather than being driven by a single transcript (**Fig. 3E**).

**Figure 3.**
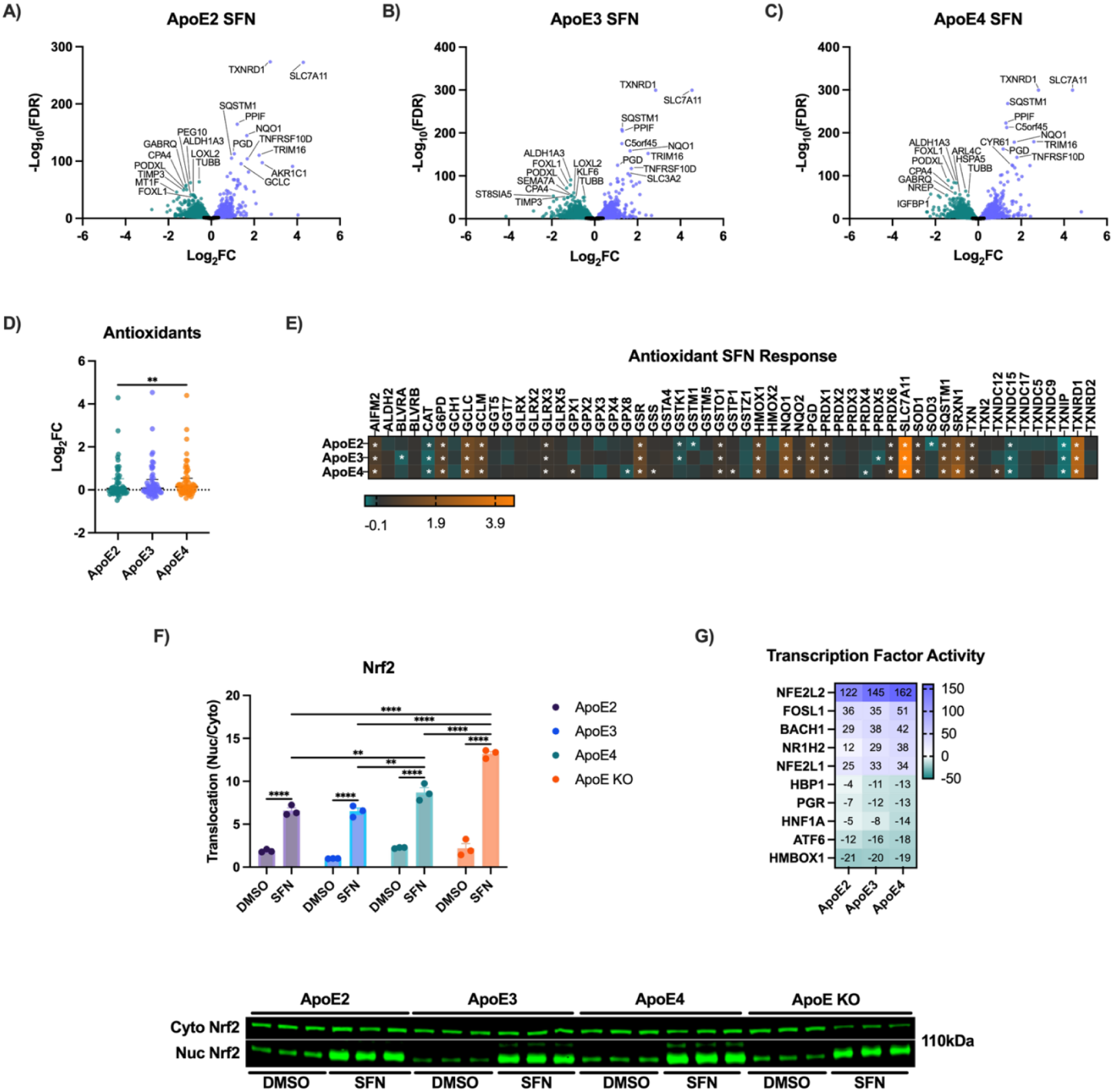
ApoE isoforms differ in their transcriptional and regulatory responses to sulforaphane. U87 cells expressing ApoE2, ApoE3 or ApoE4 were treated with DMSO or sulforaphane (SFN) for 18 hours (n=3/genotype/treatment). RNA sequencing used the ApoE-overexpression model, and protein analyses used the ApoE KO-reconstituted model, which also included ApoE KO cells (**Ext. Fig. 6**). Volcano plots of SFN-induced transcriptional responses in **A)** ApoE2, **B)** ApoE3 and **C)** ApoE4 cells. **D)** Distribution of SFN-induced log_2_ fold changes for 55 genes detected from the predefined 62-gene antioxidant panel; each point represents one gene, and the dotted line denotes no change. **E)** Heatmap of SFN-induced log_2_ fold changes for the same 55 antioxidant genes across ApoE isoforms. **F)** Nrf2 translocation following DMSO or SFN treatment in ApoE2, ApoE3, ApoE4 and ApoE KO cells; representative immunoblots are shown below. **G)** DoRothEA transcription factor activity scores for regulators associated with the SFN response across ApoE isoforms. Protein data are shown as mean±SEM with individual values overlaid. A-C) by DESeq2 Wald test with Benjamini-Hochberg correction. D) by Friedman test across isoforms (*P*=0.0077) and two-sided Wilcoxon signed-rank test against zero (ApoE2, *P*=0.058; ApoE3, *P*=0.0081; ApoE4, *P*=0.0003). F) by two-way ANOVA with ApoE genotype and treatment as factors followed by Tukey’s multiple-comparisons test. G) by permutation test with Benjamini-Hochberg correction. *p<0.05, **p<0.01, ***p<0.001, ****p<0.0001.

To determine whether this transcriptional gradient reflected upstream Nrf2 activation, we measured nuclear Nrf2 and inferred transcription-factor activity. SFN increased nuclear Nrf2 in every ApoE background and in ApoE KO cells. ApoE4 had the highest basal and post-treatment mean nuclear Nrf2 abundance among the ApoE-expressing lines, whereas ApoE3 showed the largest relative increase from baseline (**Fig. 3F**). DoRothEA independently inferred stronger NFE2L2/Nrf2 and BACH1 activity toward ApoE4 (**Fig. 3G**).

The SFN x ApoE ordinal interaction identified 67 significant genes: 43 negative coefficients indicated stronger responses toward ApoE4 and 24 positive coefficients stronger responses toward ApoE2 (**Ext. Fig. 7E**). SQSTM1 and FTH1 were among the ten most SFN-responsive genes in every isoform and had negative interaction coefficients of -0.212 and -0.187, respectively, demonstrating further amplification toward ApoE4 of transcripts central to proteostasis and iron handling. Interaction GSEA identified protein-secretion and hypoxia responses toward ApoE2 and p53 and MYC-target responses toward ApoE4 (**Ext. Fig. 7F**). BACH1, which counterbalances Nrf2-dependent transcription, also differed by isoform: SFN reduced BACH1 by 72% in ApoE2 but increased it by 173% and 161% in ApoE3 and ApoE4, respectively (**Ext. Fig. 7G**). These findings established that ApoE4 mounts a strong Nrf2-dependent transcriptional response, shifting the focus to antioxidant protein production.

### ApoE4 uncouples Nrf2-responsive transcription from antioxidant protein output

ApoE4 was associated with lower antioxidant proteins *in vivo* but mounted the strongest antioxidant RNA response to SFN. We therefore examined whether transcriptional induction produced corresponding protein accumulation, focusing on SQSTM1, GCLC and FSP1, which support proteostasis, glutathione synthesis and GPx4-independent ferroptosis suppression, respectively. ApoE KO provided a loss-of-ApoE comparator (**Fig. 4A**).

**Figure 4.**
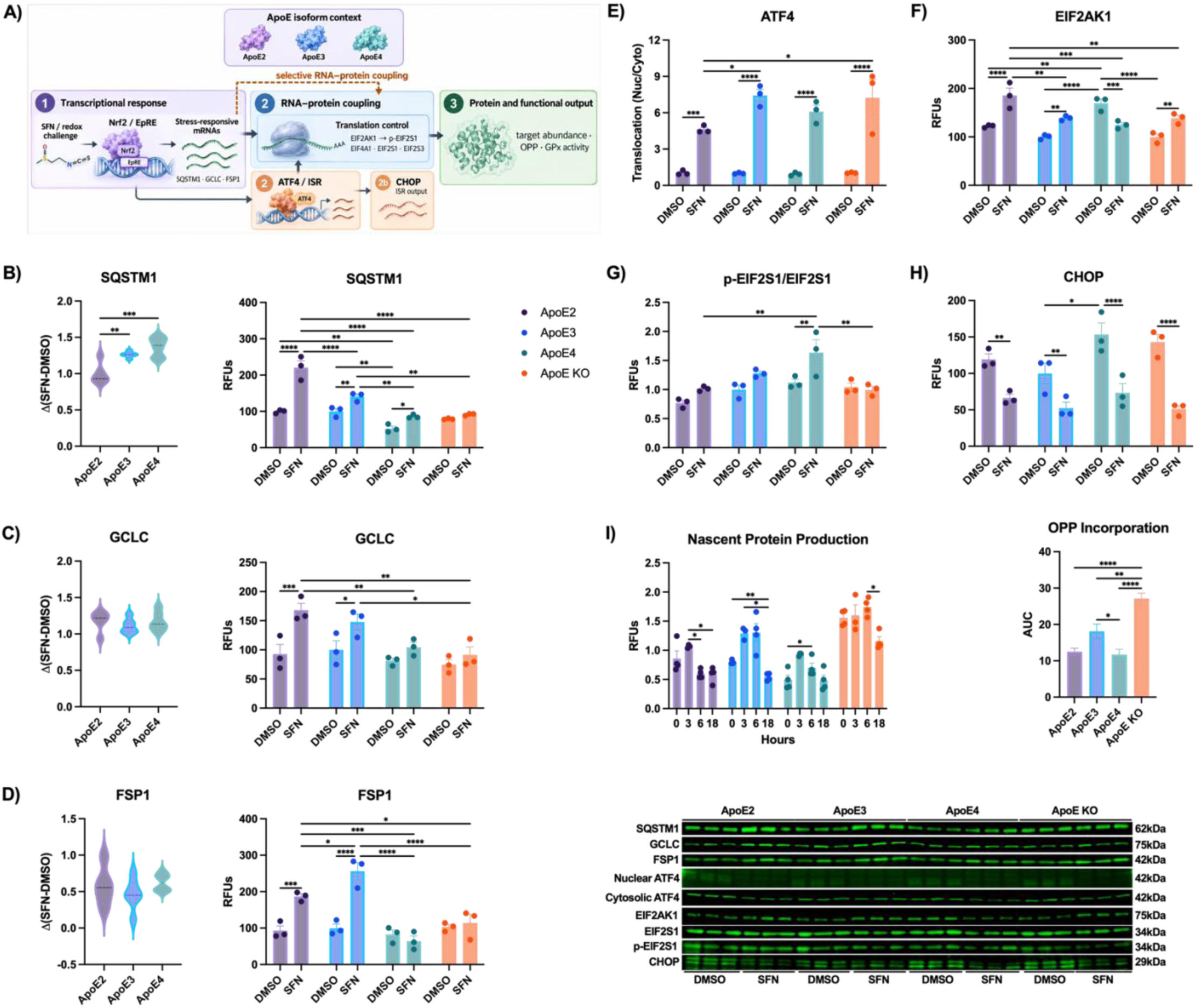
ApoE isoforms exhibit selective RNA-protein discordance following sulforaphane treatment. U87 cells expressing ApoE2, ApoE3 or ApoE4 were treated with DMSO or SFN for 18 hours (n=3/genotype/treatment); ApoE KO cells were included as a comparator for protein and OPP analyses. RNA changes were measured in the ApoE-overexpression model, and protein and OPP measurements in the ApoE KO-reconstituted model (Extended Data Fig. 6). **A)** Conceptual schematic illustrating the relationship between SFN-induced Nrf2/EpRE transcription, ATF4/integrated stress response signaling, translation control and protein/functional output; the dashed connector denotes selective RNA-protein coupling. SFN-induced RNA change [Δ(SFN–DMSO)] and corresponding protein abundance for **B)** SQSTM1, **C)** GCLC and **D)** FSP1. Integrated stress response and translation-associated immunoblots for **E)** ATF4 translocation, **F)** EIF2AK1, **G)** p-EIF2S1 (Ser51)/EIF2S1 ratio and **H)** CHOP. **I)** Nascent protein synthesis measured by O-propargyl-puromycin (OPP) incorporation at 0, 3, 6 and 18 hours after SFN treatment, with cumulative incorporation summarized as area under the curve (AUC) over 18 hours. Representative immunoblots are shown below the corresponding quantification. Data are shown as mean±SEM with individual measurements where shown. B-D) ΔRNA by one-way ANOVA with Tukey’s multiple-comparisons test. B-H) protein by two-way ANOVA with ApoE genotype and treatment as factors followed by Tukey’s multiple-comparisons test. I) time course by mixed-effects model with time, ApoE genotype and their interaction followed by Tukey’s multiple-comparisons test; AUC by one-way ANOVA with Tukey’s multiple-comparisons test. *p<0.05, **p<0.01, ***p<0.001, ****p<0.0001.

Although all three transcripts were induced, post-treatment protein abundance differed markedly. SQSTM1 was highest in ApoE2, intermediate in ApoE3 and lowest in ApoE4, which resembled ApoE KO. Despite increasing by 62% relative to DMSO, ApoE4 accumulated less SQSTM1 protein than the other isoforms while having the highest transcript abundance (**Fig. 4B**). GCLC increased by 81% in ApoE2 and 48% in ApoE3 but not significantly in ApoE4 or ApoE KO, which remained the lowest post-SFN groups (**Fig. 4C**). FSP1 doubled in ApoE2 and increased by 156% in ApoE3 but did not increase significantly in ApoE4, again resembling ApoE KO (**Fig. 4D**). Normalized RNA abundance confirmed induction of these targets, while matched analyses of FTH1, FTL, SLC7A11, HMOX1, NQO1 and GSR demonstrated that reduced protein accumulation was target-selective (**Ext. Fig. 8A-I**). Functional responses also differed: SFN increased PCOOH activity in ApoE2 and ApoE3, whereas ApoE4 retained its higher basal activity; total GPx activity did not increase in any background (**Ext. Fig. 8J,K**).

To examine regulation between RNA induction and protein accumulation, we assessed the integrated stress response. Phosphorylation of EIF2S1/eIF2α restrains general translation initiation while favoring ATF4 translation and downstream CHOP signaling. SFN increased nuclear ATF4 in every ApoE background (**Fig. 4E**). EIF2AK1, an upstream eIF2α kinase, increased in ApoE2, ApoE3 and ApoE KO but decreased in ApoE4 ( **Fig. 4F**). Nevertheless, the p-EIF2S1/EIF2S1 ratio increased selectively in ApoE4 (**Fig. 4G**). CHOP was elevated basally in ApoE4 relative to ApoE3 and decreased after SFN across backgrounds (**Fig. 4H**). Nascent protein synthesis was then measured by O-propargyl-puromycin incorporation. ApoE4 incorporation increased transiently at 6 hours and returned toward baseline at 18 hours, whereas ApoE3 incorporation decreased at 18 hours and ApoE KO produced the largest cumulative signal. The 0-18-hour area under the curve was lower in ApoE4 than ApoE3, indicating altered synthesis kinetics rather than complete global suppression (**Fig. 4I**).

We next examined NFE2L2 promoter usage, RNA processing and translation-initiation machinery as potential sources of RNA-protein discordance (**Ext. Fig. 9A**). NFE2L2 P1/P2 transcripts differ in their first exons, untranslated regions and translational efficiencies, but their relative usage showed no ApoE-dependent SFN response (**Ext. Fig. 9B**). Antioxidant-gene intronic fractions likewise showed no coordinated isoform difference, and their interaction estimates were not associated with the corresponding RNA-response interactions (**Ext. Fig. 9C,D**). EISA compared exonic and intronic SFN responses across 51 antioxidant genes to distinguish transcription-associated changes from a post-transcriptional component; this component did not differ systematically across isoforms (**Ext. Fig. 9E,F**). EIF4A1 and EIF2S3, which support mRNA scanning and eIF2-mediated initiation, respectively, also showed isoform-specific changes (**Ext. Fig. 9G,H**). Thus, reduced accumulation of selected antioxidant proteins accompanied altered eIF2 signaling and translation kinetics, without a corresponding panel-wide change in the RNA-processing measures examined.

### APOE genotype establishes region- and cell-type-dependent antioxidant states in human brain

Finally, we asked whether ApoE-dependent antioxidant states were evident before dementia and differed between vulnerable association cortex and relatively spared cerebellum. We analyzed both regions from predominantly case-matched cognitively normal older adults carrying APOE ε3/ε3 or ε3/ε4 (**Fig. 5A**). Cortex contained 430 DEGs, including ε3/ε3-biased HMOX1 and NQO1 (**Fig. 5B**). Cerebellum contained 1,929 DEGs and a broader antioxidant signature encompassing SQSTM1, GCLC and GCLM (**Fig. 5C**).

**Figure 5.**
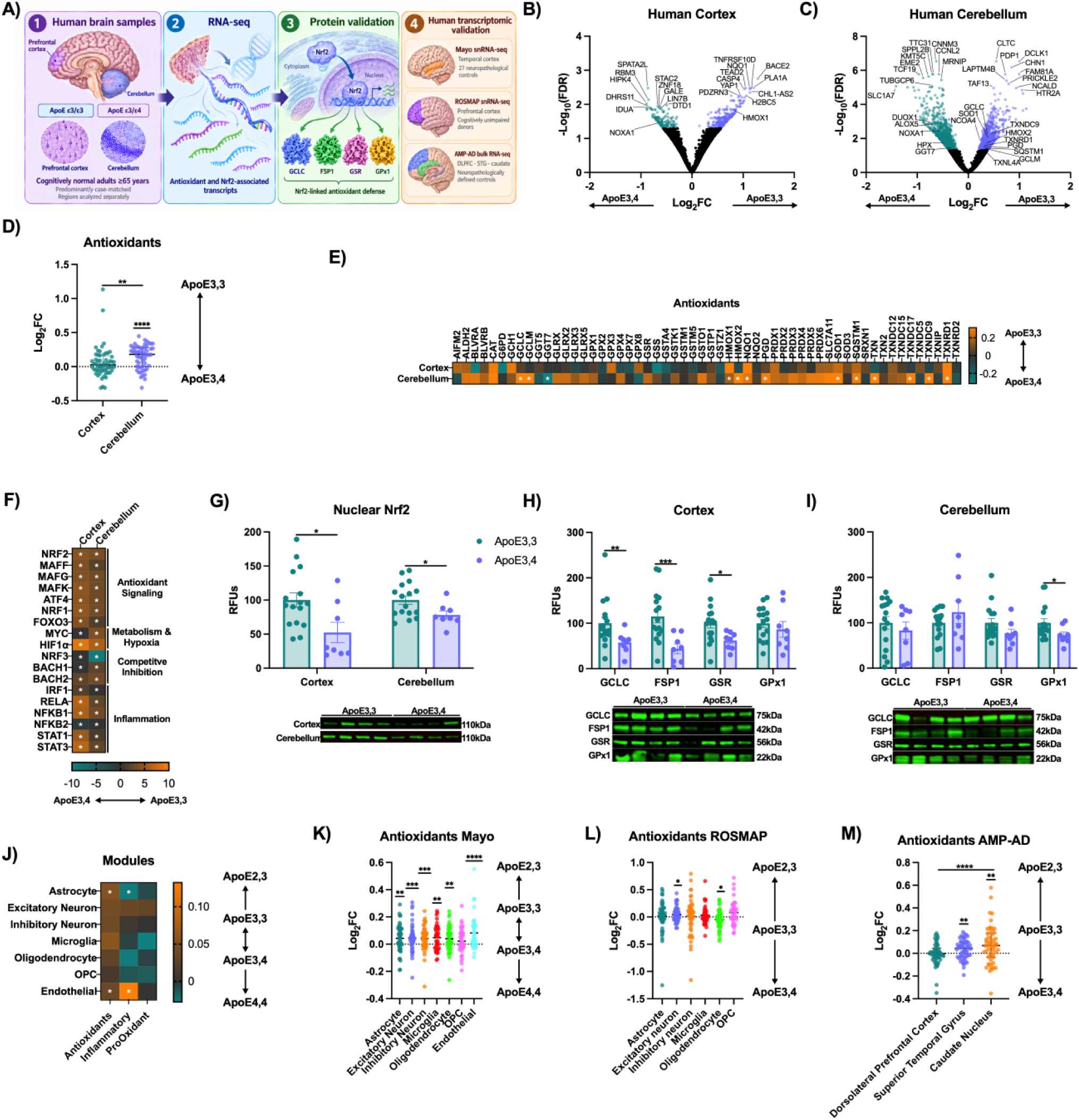
APOE genotype establishes region- and cell-type-dependent antioxidant states in the human brain. **A)** Study framework integrating regional RNA sequencing and antioxidant protein measurements from predominantly case-matched cognitively normal APOE ε3/ε3 and ε3/ε4 adults with human transcriptomic analyses from Mayo temporal cortex, ROSMAP prefrontal cortex and the AMP-AD Diverse Cohorts study. DEGs between APOE ε3/ε3 and ε3/ε4 carriers in **B)** cortex and **C)** cerebellum; positive log_2_ fold changes indicate higher expression in ε3/ε3 carriers. **D)** Log_2_ fold-change distributions for antioxidant genes detected in both regions. **E)** Heatmap of regional antioxidant-gene log_2_ fold changes. **F)** DoRothEA-inferred activity of Nrf2 and associated stress-response transcription factors. **G)** Nuclear Nrf2 abundance in cortex and cerebellum. Protein abundance of GCLC, FSP1, GSR and GPx1 in **H)** cortex and **I)** cerebellum. **J)** Heatmap of median ordered-genotype coefficients for antioxidant, inflammatory and pro-oxidant panels across major cell populations from Mayo temporal-cortex snRNA-seq of 27 control donors (APOE ε2/ε3, n=9; ε3/ε3, n=10; ε3/ε4, n=5; ε4/ε4, n=3). Genotypes were ordered ε4/ε4→ε3/ε4→ε3/ε3→ε2/ε3; positive coefficients indicate increasing expression toward ε2/ε3 and negative coefficients toward ε4/ε4. **K)** Gene-level ordered-genotype coefficients for the antioxidant panel across the indicated Mayo cell populations. **L)** Antioxidant-panel ordinal coefficients from donor-level pseudobulk analysis of ROSMAP prefrontal-cortex snRNA-seq in cognitively unimpaired donors (n=133–143, depending on cell type). **M)** Antioxidant-panel ordinal coefficients from AMP-AD Diverse Cohorts bulk RNA-seq in neuropathologically defined controls: dorsolateral prefrontal cortex (n=124), superior temporal gyrus (n=149) and caudate nucleus (n=72). For L,M, ordinal coding was ε3/ε4=−1, ε3/ε3=0 and ε2/ε3=+1; positive coefficients indicate an ε2/ε3 bias and negative coefficients an ε3/ε4 bias. Distribution plots show individual genes with median and interquartile range; protein data show individual measurements with mean±SEM. B,C,E) DESeq2 Wald tests with Benjamini-Hochberg correction. D) two-sided Wilcoxon signed-rank tests against zero within regions and a Wilcoxon matched-pairs signed-rank test between regions. F) permutation tests with Benjamini– Hochberg correction. G-I) two-tailed unpaired t-tests or Mann-Whitney tests, as appropriate. J) CAMERA-PR with Benjamini-Hochberg correction. K-M) two-sided Wilcoxon signed-rank tests against zero; regional comparisons in M by Friedman test followed by Dunn’s multiple-comparisons test. Gene-level differential expression in J-L was modeled using donor-level pseudobulk edgeR and in M using DESeq2. *p<0.05, **p<0.01, ***p<0.001, ****p<0.0001.

Across 56 shared antioxidant genes, median ε3/ε3-versus-ε3/ε4 log_2_ fold changes were +0.025 in cortex and +0.179 in cerebellum. The cerebellar distribution differed from zero, whereas the cortical distribution did not, and the paired regional distributions differed with a higher median for cerebellum (**Fig. 5D**). The heatmap showed that the cerebellar bias extended across multiple genes (**Fig. 5E**). GSEA identified ε3/ε3-biased TNFα/NF-κB and inflammatory responses in cortex and reactive oxygen species and unfolded-protein responses in cerebellum. Inflammatory distributions differed between regions, whereas pro-oxidant genes showed no coordinated genotype bias (**Ext. Fig. 10A-D**). Antioxidant genes were enriched among ε3/ε3-biased cerebellar DEGs (**Ext. Fig. 10E**).

We then examined whether these RNA differences were accompanied by altered antioxidant regulation and protein abundance. DoRothEA inferred an ε3/ε3 bias in Nrf2 activity in both regions, together with its small-Maf partners and associated stress-responsive transcription factors (**Fig. 5F**). Nuclear Nrf2 was 48% lower in ε3/ε4 cortex and 22% lower in cerebellum (**Fig. 5G**). Cortical GCLC, FSP1 and GSR were 28%, 62% and 35% lower, respectively, while GPx1 was unchanged (**Fig. 5H**). Cerebellar GCLC, FSP1 and GSR were preserved, whereas GPx1 was lower (**Fig. 5I**). The region with the weaker coordinated RNA difference therefore carried the broader protein deficit.

To resolve cellular contributions, we analyzed temporal-cortex snRNA-seq from 27 donors without AD neuropathology spanning APOE ε2/ε3, ε3/ε3, ε3/ε4 and ε4/ε4 from the Mayo Clinic Brain Bank cohort^20^. Genotypes were ordered ε4/ε4→ε3/ε4→ε3/ε3→ε2/ε3. Antioxidant coefficients favored ε2/ε3 across major neural and vascular populations, with significant competitive panel associations in astrocytes and endothelial cells ( **Fig. 5J,K**). We tested reproducibility in two larger cohorts. Donor-level pseudobulk analysis of ROSMAP prefrontal-cortex snRNA-seq from cognitively unimpaired individuals showed an ε2/ε3-biased antioxidant state most clearly in excitatory and inhibitory neurons, with median ordinal coefficients of +0.040 and +0.058, respectively ( **Fig. 5L**). In AMP-AD neuropathologically defined controls, bulk antioxidant expression favored ε2/ε3 in superior temporal gyrus and caudate nucleus, with median coefficients of +0.044 and +0.068, but showed little overall bias in dorsolateral prefrontal cortex (**Fig. 5M**).

These antioxidant relationships differed from inflammatory regulation. In Mayo temporal cortex, inflammatory coefficients favored ε4 in astrocytes and oligodendrocytes but ε2/ε3 in excitatory neurons and endothelial cells; pro-oxidant expression favored ε2/ε3 only in excitatory neurons (**Ext. Fig. 10F,G**). ROSMAP similarly showed ε3/ε4-biased inflammatory expression in astrocytes, oligodendrocytes and OPCs but an ε2/ε3 bias in excitatory neurons, without a coordinated pro-oxidant shift (**Ext. Fig. 10H,I**). In AMP-AD, inflammatory expression favored ε3/ε4 in dorsolateral prefrontal cortex and superior temporal gyrus, with the strongest bias in dorsolateral prefrontal cortex (**Ext. Fig. 10J**). A pro-oxidant bias toward ε3/ε4 was detected only in dorsolateral prefrontal cortex (**Ext. Fig. 10K**). Across three independent cohorts, older human brain therefore exhibited an ApoE2-associated antioxidant transcriptional state whose expression depended on region and cell type. Our regional protein measurements further showed that the magnitude of this transcriptional difference did not predict the antioxidant proteins available in cognitively normal brain.

## Discussion

ApoE4 is most notably associated with increased risk for Alzheimer’s disease. Our findings identify differences in stress adaptation that precede clinical disease and change with age and cellular context. Young ApoE4 brains had lower antioxidant protein abundance and reduced oxidized phospholipid detoxification before detectable oxidative damage, whereas aged ApoE4 brain accumulated lipid peroxidation despite antioxidant remodeling and higher nuclear Nrf2. Acute SFN stimulation exposed a related separation: ApoE4 cells mounted the strongest antioxidant transcriptional response but accumulated less protein from selected targets. Antioxidant regulation therefore depends on how transcriptional activation is coupled to protein accumulation and functional defense.

The proteins examined converge on protection against lipid peroxidation. GCLC and GCLM sustain glutathione synthesis^23^, supporting reduction of oxidized phospholipids by GPx4 and Prdx6^24–27^, while FSP1 maintains reduced coenzyme Q as a parallel defense against lipid-radical propagation^28,29^. Reduced protein abundance, together with lower PCOOH activity, defines diminished ferroptosis-suppressive reserve in young ApoE4 brain. The later increase in HNE shows that antioxidant remodeling does not preserve redox homeostasis indefinitely. This connects ApoE-dependent stress adaptation to the iron-associated lipid damage documented in AD and Down syndrome with AD^10,11^. Evidence that protective ApoE variants support ABCA7-dependent efflux of oxidized neuronal phospholipids further links ApoE biology to ferroptotic regulation^12^.

The aging analysis places these biochemical findings within an adaptive transcriptional landscape. Antioxidant expression favored ApoE4 in young mice and isogenic human neuron-astrocyte co-cultures but shifted toward ApoE2 in aged mouse cortex. Elevated antioxidant RNA in young ApoE4 systems is therefore compatible with greater compensatory demand despite lower protein defense. Independent pathway enrichment and the external Nrf2 regulon supported this transition beyond the predefined antioxidant panel. The opposing pro-oxidant trajectory and less uniform inflammatory response further indicate selective remodeling of antioxidant maintenance rather than a common shift across global stress-associated genes. Elevated basal antioxidant abundance can coexist with reduced inducibility: older mice had higher nuclear Nrf2 and GCLC/GCLM levels but showed little further induction following chronic ambient nanoparticle exposure^30^. ApoE-dependent antioxidant abundance must consequently be interpreted in relation to age and regulatory context.

Adult allelic switching showed that these transcriptional states remain modifiable. Global and astrocyte-restricted APOE4-to-APOE2 switching redirected antioxidant expression in glia, including microglia whose genotype remained unchanged after astrocyte-restricted switching. This identifies an astrocyte-dependent contribution to antioxidant regulation across cell types, extending established cell-autonomous and non-cell-autonomous effects of ApoE^21^. The switched state also differed from constitutive young ApoE2 mice, suggesting that current isoform expression and prior cellular history jointly influence the response. The metabolic, pathological and cognitive improvements reported in the switching model place this plasticity within a broader functional context^22^.

The expanded human analyses extend the late-life ApoE2-associated antioxidant state across Mayo, ROSMAP and AMP-AD datasets^20,31,32^. Its distribution differed: Mayo temporal cortex showed broad cellular involvement, ROSMAP prefrontal cortex showed clearer neuronal associations, and AMP-AD bulk tissue showed stronger antioxidant biases in superior temporal gyrus and caudate than dorsolateral prefrontal cortex. Inflammatory and pro-oxidant relationships followed different cellular and regional patterns. These findings explain why a single bulk-tissue comparison cannot fully describe ApoE-dependent stress regulation. Our regional protein measurements exposed an additional separation: cerebellum showed the stronger antioxidant RNA difference while preserving most measured proteins, whereas cortex exhibited a smaller coordinated RNA shift but broader reductions in GCLC, FSP1 and GSR, which parallel the associated AD neuropathology found in these brain regions. Transcriptional differences therefore provide only part of the explanation for regional antioxidant capacity.

Our SFN experiments connect ApoE-dependent antioxidant regulation to stress-responsive control of protein synthesis. ApoE4 cells retained nuclear Nrf2 accumulation and ATF4 induction alongside increased eIF2α phosphorylation and altered nascent-protein-synthesis kinetics. Increased eIF2α phosphorylation is already evident in young ApoE4 mice^13^, while primary ApoE4 microglia exhibit increased phosphorylation of eIF2α and its upstream kinases together with reduced basal protein synthesis^33^. Microglial ISR activation also promotes secretion of neurotoxic long-chain lipids, and inhibiting either ISR signaling or lipid synthesis mitigates synapse loss in AD models^34^. Translation recovery requires relief of eIF2B inhibition: PPP1R15B enables dephosphorylation of eIF2 within the inhibited eIF2-eIF2B complex, restoring translation initiation^35^. Proteasome inhibition also sensitizes cells to GPx4 inhibition through a mechanism requiring new protein synthesis, while ATF4 induction opposes this sensitization, linking protein turnover and stress-responsive synthesis to ferroptosis susceptibility^36^. Isogenic human neuronal analyses likewise report ApoE4-associated changes in ribosome occupancy and codon-specific pausing alongside slower protein renewal^37^. These findings connect the activation and resolution of stress signaling to protein production and lipid toxicity, providing a framework for the selective antioxidant RNA-protein discordance observed here. NFE2L2 promoter usage, intronic fractions and EISA showed no coordinated isoform effect accounting for that discordance, directing attention toward transcript-specific regulation, translation efficiency and protein turnover.

The ancestral status of ApoE4 supports an evolution-informed interpretation of these stress-adaptation states. ApoE4 retains the ancestral residues at positions 112 and 158, whereas ApoE3 and ApoE2 are successively derived isoforms^3,38^. ApoE3 midpoint centering among genes with intermediate expression supports comparing transcriptional states along this sequence. APOE2-associated longevity persists independently of AD status, and median survival in targeted-replacement mice follows ApoE2>ApoE3>ApoE4^2^. Human APOE2 neurons also show enhanced DNA-repair signaling and resistance to stress-induced DNA damage and senescence^14^. Proteomic analyses similarly associate APOE2 with DNA repair, translation and autophagy, with many differences already detectable in amyloid-negative individuals^8^. These maintenance-associated effects are cell-type dependent: plasma-proteomic aging estimates indicate younger astrocyte but older macrophage profiles in APOE2 carriers, with the opposite pattern in APOE4 carriers^39^. Our age-dependent antioxidant reversal and distinct neuronal, glial and regional associations fit this broader organization, in which ApoE isoforms favor different forms of cellular maintenance according to age and biological context.

An evolutionary trade-off between infectious defense and long-term tissue preservation offers a framework for this organization. APOE ε4 is associated with higher fecundity and better cognitive performance under high parasite burden^5,6^, whereas chronic low-grade inflammation increases AD risk disproportionately among ε4 carriers^15^. These relationships indicate that the consequences of an ApoE-associated immune state depend on environmental conditions^38,40^. Nrf2 connects antioxidant maintenance with inflammatory regulation by inducing protective genes while suppressing inflammatory cytokine transcription^16^. Our results fit a model in which the derived isoforms increasingly support antioxidant maintenance during prolonged sterile inflammation, while the ancestral state remains advantageous in other settings. Declining metabolic reserve with age would make differences in sustained protein defense increasingly consequential.

The inverse APOE relationship in melanoma illustrates the same contextual dependence. APOE2 is associated with more aggressive melanoma metastasis, whereas APOE4 is protective^7^, and ApoE2 stimulates protein synthesis through LRP1 in melanoma cells^41^. Malignant cells also exploit Nrf2-dependent antioxidant and ferroptosis-suppressive defenses to survive metabolic stress^42,43^. The capacity to sustain protein production and resist lipid peroxidation can therefore support either tissue maintenance or malignant-cell survival, depending on cellular context. This places the antioxidant phenotype within a broader relationship between ApoE genotype, stress tolerance and disease susceptibility.

Several limitations are present. RNA and protein responses were measured in independently generated U87 systems, and OPP measured global synthesis rather than translation of individual targets. Human analyses were cross-sectional and used cognitive or neuropathological inclusion criteria, limiting inference about individual aging trajectories. The findings establish selective differences in antioxidant regulation and protein accumulation; the molecular mechanism connecting them remains unresolved.

These findings have implications for AD trials of sulforaphane (NCT04213391), dimethyl fumarate (NCT06850597) and hydralazine (NCT04842552)^44^ which enhance Nrf2 by acting on its cytosolic repressor protein Keap1. The stronger antioxidant transcriptional response in ApoE4 cells yielded less protein from selected protective targets, identifying APOE genotype as a potential determinant of pharmacodynamic response. Prespecified APOE-by-treatment analyses, together with protein and functional measurements, would therefore help determine whether transcriptional target engagement translates into antioxidant protection.

ApoE-associated antioxidant vulnerability thus emerges as a dynamic, partially reprogrammable property of aging. Reduced phospholipid-hydroperoxide detoxification precedes detectable damage, while strong stress-responsive transcription does not ensure accumulation of key proteins supporting proteostasis, glutathione synthesis and ferroptotic suppression. These findings provide critical insight for how APOE ε4 may accelerate AD onset and progression: reduced antioxidant reserve and inadequate protective protein accumulation create a mismatch between stress-response activation and effective defense, allowing lipid injury to accumulate and progressively erode brain resilience with age.

## Methods

### Mice

All animal procedures were approved by the University of Southern California Institutional Animal Care and Use Committee under protocols 20842, 11678 and 11233.Two humanized ApoE targeted-replacement mouse models expressing human ApoE3 or ApoE4 were used. The first cohort comprised 5xFAD-negative mice from the EFAD line (referred to here as ApoE-TR), which was generated by crossing ApoE targeted-replacement mice with 5xFAD mice on a 129xC57BL/6 background and subsequently backcrossing to C57BL/6. Mice that did not inherit the 5xFAD transgenes express human ApoE3 or ApoE4 from the endogenous murine locus without familial AD transgenes and were provided by Mary Jo LaDu and Leon Tai.

The second model (CAF-TR) was generated by the Cure Alzheimer’s Fund and Holtzman laboratory by replacing the murine Apoe coding sequence with human APOE3 or APOE4 on a C57BL/6 background. The targeted allele contains loxP-flanked sequences that permit conditional disruption and was generated without a neomycin-selection cassette^45^. Both models express human ApoE under endogenous regulatory control. ApoE-TR and CAF-TR mice were studied at 6 months; CAF-TR mice were also studied at 22 months. Both sexes were included for all experiments. Exact numbers for each genotype, age, tissue and assay are reported in the figure legends and source data.

### Tissue collection

Mice were anesthetized with isoflurane, euthanized by cardiac puncture and transcardially perfused with PBS. Brains were hemisected and dissected into cortex, hippocampus and cerebellum. Lung, liver, heart and midbrain subregions were collected from ApoE-TR mice for Extended Data analyses. Tissues were snap-frozen on dry ice and stored at −80 °C.

### Primary mixed glia

Primary mixed glia were generated from cerebral cortices of postnatal day 3-4 CAF-TR mice. Cortical tissue was dissociated and maintained in DMEM/F12 media (Gibco, ThermoFisher Scientific, Waltham, MA) supplemented with 10% fetal bovine serum, 1% penicillin, and 1% L-glutamine (Gibco, ThermoFisher Scientific, Waltham, MA) at 37°C, 5% CO ^46,47^. Cells were harvested once 100% confluency was reached and processed for protein extraction.

### U87 ApoE cell lines and sulforaphane treatment

U87 glioblastoma and HEK293T cells were obtained from ATCC. Cells were maintained in DMEM supplemented with 10% fetal bovine serum and 1% penicillin-streptomycin at 37°C in a humidified incubator containing 5% CO2. For dose selection, U87 cells were treated for 18 h with 0, 2.5, 5, 7.5 or 10μM sulforaphane (SFN), and the expression of Nrf2-responsive genes and proteins was evaluated by quantitative PCR and immunoblotting. A concentration of 7.5μM produced robust induction across the evaluated endpoints and was selected for subsequent experiments. DMSO was used as the vehicle control.

### Generation of APOE-expressing U87 models

Two independently generated U87 models were used and analyzed separately. The first model, used for the SFN transcriptomic experiments, expressed APOE2, APOE3 or APOE4 on the endogenous APOE3/3 U87 background. APOE isoforms were introduced using UBC promoter-driven lentiviral constructs that co-expressed an mCherry reporter (Addgene plasmids 190807, 190808 and 190809 for APOE2, APOE3 and APOE4, respectively).

For subsequent protein and functional experiments, endogenous APOE was disrupted before isoform reconstitution. U87 cells were first transduced with lentiviral particles encoding Cas9 nuclease and selected with puromycin to establish a stable Cas9-expressing population. Cells were then transfected with four guide RNAs targeting APOE using Lipofectamine RNAiMAX. The guide sequences were: CCTGGTGCAGTACCGCGGCG, GGCCTACAAATCGGAACTGG, CTTTTGGGATTACCTGCGCT and GCTGATGGACGAGACCATGA. Cells were expanded 48h after transfection, and APOE disruption was evaluated by genotyping and quantitative PCR, with loss of ApoE protein confirmed by immunoblotting.

APOE2, APOE3 and APOE4 open reading frames were cloned into the pReceiver-Lv183 lentiviral backbone using HiFi DNA assembly and introduced separately into the APOE-knockout cells. Six independent transductions were generated for each APOE isoform and served as biological replicates. An unreconstituted APOE-knockout population was retained as a loss-of-ApoE comparator. The two generations of U87 models were not pooled for analysis.

### Lentivirus production and transduction

HEK293T cells were seeded at approximately 40% confluence and transfected with the appropriate expression construct using a lentiviral packaging system in antibiotic-free medium. Medium was replaced after 18 h, and lentivirus-containing medium was collected 24 and 48 h later. Viral preparations were passed through a 0.45-μm polyethersulfone filter and concentrated through a sucrose gradient by ultracentrifugation at 70,000g for 2.5h at 4°C. Viral pellets were resuspended in 10μl ice-cold DPBS per 10 ml of collected viral medium.

U87 cells were exposed to concentrated lentiviral particles at a 1:500 dilution in antibiotic-free medium containing 8 μg/mL polybrene. After 18-24 h, the transduction medium was replaced with complete medium. Antibiotic selection was initiated 48 h after transduction using 3.5μg/ml puromycin or 2mg/ml neomycin, according to the resistance cassette carried by the corresponding construct.

### Cell-line and construct verification

APOE expression constructs were sequence-verified before lentiviral production. Following transduction, APOE isoform identity was reconfirmed by sequencing APOE amplified from the corresponding cell populations. These sequencing procedures verified the introduced APOE construct and isoform but were not used as a substitute for short tandem repeat profiling of the parental cell lines. APOE expression was additionally assessed by quantitative PCR and immunoblotting. Depletion of ApoE in the knockout model was independently supported by mass spectrometry. Cell model validation details are in **Extended Figure 6**. Routine mycoplasma screening was performed every six months.

### Human postmortem brain tissue

Postmortem prefrontal cortex and cerebellum were obtained from the University of Southern California Alzheimer’s Disease Research Center and University of California, Irvine MIND program under USC IRB protocol UP-20-00014-EXEMPT and the consent procedures of the contributing brain banks. Samples were from cognitively normal adults aged 65 years or older carrying APOE ε3/ε3 or ε3/ε4. The RNA-sequencing cohort comprised 22 cortex (Brodmann Areas 8/9/10) samples (16 ε3/ε3 and 6 ε3/ε4) and 31 cerebellum samples (16 ε3/ε3 and 15 ε3/ε4); 21 donors contributed both regions. Individual characteristics are provided in **Supplementary Table 1**. APOE genotype was determined from rs429358 and rs7412.

### Subcellular fractionation and immunoblotting

Frozen tissue (20mg) was homogenized in 125 μl sucrose-based fractionation buffer for separation of nuclear and cytosolic fractions^48^. For brain tissue, 1% Triton X-100 was added to the initial wash and the resuspended nuclear pellet was incubated on ice for 30 min to lyse residual cells. Fraction purity was assessed using histone H3 and GAPDH as nuclear and cytosolic markers, respectively. The same fractionation strategy was used for U87 localization assays.

Protein concentration was determined by Bradford or bicinchoninic acid assay as appropriate. Twenty micrograms of protein was resolved on 4-15% gradient gels and transferred to 0.45-μm polyvinylidene difluoride membranes. Membranes were normalized by total-protein staining where indicated, blocked in LI-COR Intercept buffer and incubated overnight with primary antibodies. After washing in TBS containing 0.1% Tween -20, membranes were incubated with IRDye 700- and/or 800-conjugated secondary antibodies, imaged on a LI-COR Odyssey system and quantified in ImageJ.

### Dot-blot analysis of oxidative damage

Frozen tissue (20 mg) was homogenized in RIPA buffer containing protease inhibitors and centrifuged at 10,000xg for 15 min at 4°C. Equal protein amounts were applied to membranes using a Bio-Dot apparatus. HNE Michael adducts and nitrotyrosine were detected with primary antibodies, imaged as described for immunoblotting and normalized to total protein.

### Antioxidant enzyme assays

Total glutathione peroxidase and superoxide dismutase activities were measured with Cayman Chemical assays according to the manufacturer’s instructions. For phospholipid hydroperoxide-reducing activity, oxidized phosphatidylcholine hydroperoxide (PCOOH) was generated as described previously^10,11,49,50^ and substituted for the supplied cumene hydroperoxide. The same PCOOH preparation was used for all samples within an experiment. Absorbance was measured using a temperature-controlled SpectraMax M2 spectrophotometer and normalized to protein measured by the 660-nm assay.

### Nrf2 chromatin immunoprecipitation-quantitative PCR

Nrf2 promoter occupancy was measured in cortex from 6-month-old ApoE3 and ApoE4 CAF-TR mice. Tissue (100 mg) was washed and minced in PBS, crosslinked in 1% formaldehyde for 10 min at room temperature and quenched with 2.5M glycine for 5 min. After three ice-cold PBS washes, tissue was disrupted in RIPA buffer (0.5 M Tris-HCl, pH 7.4, 1.5 M NaCl, 2.5% deoxycholic acid, 10% NP-40 and 10 mM EDTA), incubated on ice for 30 min and centrifuged at 20,000xg for 15 min at 4°C. Pellets were resuspended in RIPA conta ining 1% SDS and sonicated using a Bioruptor UCD-200. Three cortices were pooled per biological replicate, with two independent pools per genotype.

One microgram sheared chromatin was immunoprecipitated for 16h with 1 μg anti-Nrf2 antibody or rabbit IgG. Immunoprecipitates were washed sequentially in low-salt, high-salt, LiCl and EDTA/Tris buffers and eluted overnight at 65°C in 1% SDS and 100 mM sodium bicarbonate. Protein and RNA were removed with proteinase K and RNase A before DNA purification. Published Nrf2 electrophile-response elements (EpREs) guided selection of GPx4 and Prdx6 promoter regions^51^. Primer sequences are provided in **Supplementary Table 2**. Enrichment was normalized to matched rabbit IgG controls.

### Nascent protein synthesis

Nascent protein synthesis was measured using the Click-iT Plus O-propargyl-puromycin (OPP) Alexa Fluor 488 assay. Cells were labeled with 20μM OPP for 30 min, fixed in 4% paraformaldehyde, permeabilized with 0.5% Triton x-100 and reacted with Alexa Fluor 488 picolyl azide. Fluorescence was imaged at 10x using an Incucyte system and normalized to confluence measured by phase imaging. Measurements were obtained 0, 3, 6 and 18 h after SFN treatment. The 0-18h area under the curve was calculated for each biological replicate by trapezoidal integration.

### RNA extraction, library preparation and primary processing

Frozen brain tissue (20 mg) was homogenized in 1 ml TRIzol using six 10 second cycles separated by 60 second cooling intervals. After addition of 300 μl chloroform, the aqueous phase was separated in heavy-gel phase-lock tubes and purified using Extracta Plus columns according to the manufacturer’s protocol.

Poly(A)-selected mRNA libraries were sequenced by Novogene as paired-end 150-bp reads on an Illumina NovaSeq platform, generating approximately 6 Gb per sample where applicable. Reads were trimmed with Trim Galore 0.6.5-1, aligned to the appropriate mouse or human reference genome using STAR 2.7.3a^52^ and assigned to genes using featureCounts from Subread 2.0.3^53^. Analyses were performed in R 4.3.2 with DESeq2 1.34.0^54^.

### Predefined gene panels

Analyses used predefined panels containing 62 antioxidant, 111 inflammatory and 25 pro-oxidant genes. Panel definitions were maintained across datasets, with species-appropriate gene symbols matched before extraction. Genes absent from a dataset or failing its expression filter were excluded rather than assigned zero values. Panel analyses retained all evaluable genes irrespective of individual differential-expression significance. Consequently, the number of represented genes differed among datasets, regions and cell populations.

### Aging ApoE-targeted-replacement mouse RNA-sequencing analysis

The aging analysis used gene-level counts from 143 cortical samples collected at 3, 12 and 24 months from ApoE2-, ApoE3- and ApoE4-targeted-replacement mice^19^. Counts were rounded to integers and genes with at least 10 reads in at least eight samples across the dataset were retained.

Age-specific categorical models included sex as a covariate. For ordinal analyses, genotype was encoded ApoE4=-1, ApoE3=0 and ApoE2=+1; positive coefficients therefore indicate increasing expression toward ApoE2 and negative coefficients indicate increasing expression toward ApoE4. Age-specific ordinal coefficients were estimated separately at 3, 12 and 24 months. The primary aging model used the 3 - and 24-month samples and the formula expression ∼ sex + age + ordinal ApoE + age x ordinal ApoE. The interaction coefficient quantifies the change in the ordinal slope from 3 to 24 months.

The predefined 62-gene antioxidant panel and predefined inflammatory and pro-oxidant modules were analyzed independently of transcriptome-wide significance. Module summaries are distributions of gene-level coefficients rather than enrichment scores selected by significance. Variance-stabilized expression of representative genes was analyzed using two-way ANOVA with age, genotype and their interaction, followed by Tukey correction. Transcriptome-wide significance was defined as Benjamini-Hochberg-adjusted *P*<0.05.

### Gene-set enrichment and transcription-factor activity

Pre-ranked GSEA^55^ v4.4.0.2 used transcriptome-wide DESeq2 statistics from the indicated contrast or interaction and the MSigDB Hallmark collections^56^. Normalized enrichment scores and FDR-adjusted values were reported. Transcription-factor activity was inferred separately from DESeq2 log_2_ fold changes using human DoRothEA regulons^57^ and restricted to confidence levels A-C and weighted-mean scoring implemented in decoupleR^58^. Genes lacking valid symbols or log_2_ fold-change estimates were excluded, and duplicate gene symbols were collapsed by averaging their log_2_ fold changes. Empirical two-sided P values were calculated from 1,000 permutations per contrast by shuffling log_2_ fold-change values among gene symbols and comparing the absolute observed activity score with the resulting null distribution. P values were corrected across transcription factors within each contrast using the Benjamini-Hochberg method.

### Independent Nrf2-regulon overlay

Independent mouse Nrf2 target sets were obtained from the published ChIP-seq and expression analysis of Malhotra et al^51^. The basal and inducible sets contained 654 and 645 genes, respectively, with 244 shared genes and 1,055 unique genes in their union. Published symbols were matched to the Zhao expression analysis. ApoE-responsive genes were defined by adjusted *P*<0.05 in either the ApoE2-versus-ApoE3 or ApoE4-versus-ApoE3 categorical contrast. Enrichment was tested against the remaining expressed genes using two-sided Fisher exact tests and Benjamini-Hochberg correction. The responsiveness analysis used all 25,563 expressed genes.

### ApoE3 midpoint-position analysis

Categorical ApoE2-versus-ApoE3 and ApoE4-versus-ApoE3 coefficients were used to test the ordinal relationship without imposing ordinal coding. For each gene with distinct ApoE2 and ApoE4 endpoint estimates, ApoE3 position was normalized to the ApoE4→ApoE2 interval: (ApoE3−ApoE4)/(ApoE2−ApoE4). ApoE4 therefore equals 0, ApoE2 equals 1 and 0.5 denotes the theoretical midpoint. Genes were classified as beyond ApoE4 (<0), intermediate (0–1) or beyond ApoE2 (>1); non-finite positions were excluded.

Genome-wide positional classes were compared across ages using paired Stuart-Maxwell tests with Holm correction for the three pairwise comparisons. Module-level class distributions used Stuart-Maxwell tests with Benjamini-Hochberg correction. Among intermediate genes, normalized positions were tested against 0.5 using two-sided Wilcoxon signed-rank tests with Benjamini-Hochberg correction. Paired age comparisons were restricted to genes intermediate at both ages.

### Cell-type composition and sensitivity analysis

Cell-type proportions were estimated from aging mouse bulk cortex using MuSiC^59^ and a Tabula Muris Senis cortical reference^60^. Downstream models considered neurons, astrocytes, microglia, oligodendrocytes and oligodendrocyte precursor cells. Endothelial and pericyte estimates were retained for quality assessment but excluded from adjustment because their estimated contributions were negligible. Composition was analyzed using linear models containing age, genotype, age x genotype and sex. For each cell-type-enriched sensitivity model, bulk counts were fit with the same age x ordinal ApoE design while standardized proportions of the other major cell types were included as covariates; the target cell-type proportion was intentionally omitted. These models test whether the interaction persists after accounting for signal attributable to other cell populations and are not direct single-cell expression measurements. A seven-versus-eight-cell reference analysis assessed sensitivity to reference composition.

### U87 ApoE isoform and SFN RNA-sequencing analysis

The final analysis included ApoE2-, ApoE3- and ApoE4-expressing samples treated with DMSO or SFN. Protein-coding and selected noncoding genes were retained; mitochondrial genes, small RNA classes, rRNA, pseudogenes and reporter features were excluded. Genes required at least 10 reads in at least three samples. Within-isoform SFN contrasts were estimated with an ApoE-by-treatment group factor. The SFN x ApoE ordinal interaction encoded ApoE4=0, ApoE3=1 and ApoE2=2 and used expression ∼ ordinal ApoE + SFN + SFN x ordinal ApoE. Positive interaction coefficients indicate stronger responses toward ApoE2 and negative coefficients indicate stronger responses toward ApoE4. No log_2_ fold-change threshold was applied. APOE was excluded from the control-gene set used for size-factor estimation so that normalization was not driven by the transgene. DESeq2 Wald tests used Benjamini**-**Hochberg correction; effect-size shrinkage was applied to pairwise contrasts where indicated. The antioxidant panel was evaluated independently of transcriptome-wide significance. Fifty-five of 62 genes had complete values across the three within-isoform contrasts. Variance-stabilized expression was used for representative genes, and SFN-induced RNA change was calculated as Δ(SFN−DMSO).

### NFE2L2 promoter-usage analysis

Promoter-specific NFE2L2 first exons were quantified from aligned reads for the same U87 samples using paired-end, reverse-stranded featureCounts^61^. P1 was defined as chr2:177264532-177264727 (196 bp) and P2 as chr2:177263403-177264131 (729 bp; GRCh38). Counts were divided by feature length, and P2 fraction was calculated as P2/(P1+P2). Length-normalized P1 and P2 signals were analyzed by quasibinomial regression containing ordinal ApoE, SFN and their interaction.

### Antioxidant exon-intron analysis

Exonic and intronic features were generated from GENCODE v44 for the 62-gene antioxidant panel and quantified separately from aligned U87 reads using paired-end, reverse-stranded featureCounts. Intronic fraction was calculated as intronic reads/(intronic+exonic reads). Architecture was available for 60 genes; 57 had complete values for isoform comparisons and 55 for comparison with RNA SFN x ApoE ordinal interaction estimates. Intronic and exonic counts were modeled by quasibinomial regression as intronic/exonic reads ∼ SFN x ordinal ApoE. The association between RNA and intronic-fraction interaction coefficients was tested by two-sided Spearman correlation.

### Exon-intron split analysis

Exon-intron split analysis (EISA)^62^ was used to separate transcription-associated and post-transcriptional components of the SFN response across the predefined antioxidant panel. The analysis used the exonic and intronic count matrices described above and the same U87 cohort used for the transcriptomic analyses. Whole-transcriptome DESeq2 size factors from the original SFN ordinal analysis were retained, without renormalizing the antioxidant panel independently. Genes were included in the primary EISA if they had at least 10 exonic reads in at least six samples, at least five intronic reads in at least six samples, and nonzero exonic and intronic counts summed within each of the six ApoE-by-treatment groups.

SFN-induced changes in exonic and intronic abundance were first estimated separately using negative-binomial DESeq2 models with an ApoE-by-treatment group factor, from which SFN-versus-DMSO contrasts were extracted within each ApoE isoform. The EISA post-transcriptional component (PTc) was defined as the SFN-induced change in exonic abundance relative to the corresponding change in intronic abundance: *PTc = Δexonic response − Δintronic response.* Formal PTc estimates and gene-level p-values were obtained using quasibinomial exon-to-intron ratio models. To test whether PTc changed progressively across isoforms, ApoE was encoded as ApoE4=0, ApoE3=1 and ApoE2=2, and models included SFN, ordinal ApoE and their interaction.

### Human brain RNA-sequencing analysis

Cortex and cerebellum were analyzed separately and genes with at least 10 reads in at least two samples were retained. No sample was excluded solely on the basis of PCA or sample-distance visualization.

Surrogate-variable analysis^63^ was performed separately by region on variance-stabilized expression. Three surrogate variables were included for cortex and five for cerebellum. Final DESeq2 models included sex, postmortem interval, APOE genotype and region-specific surrogate variables; age, Braak stage, RIN and cell-type deconvolution estimates were not included. The primary contrast was APOE ε3/ε4 versus ε3/ε3. Positive original coefficients denote higher expression in ε3/ε4.

Predefined antioxidant, inflammatory and pro-oxidant modules were analyzed from gene-level coefficients. Distributions were tested against zero by two-sided Wilcoxon signed-rank tests, and shared genes were compared between regions by Wilcoxon matched-pairs signed-rank tests. Directional enrichment among FDR-significant genes was tested by two-sided Fisher exact tests against the expressed-gene background, with Benjamini-Hochberg correction.

### Independent analysis of isogenic human iPSC-derived neuron–astrocyte co-cultures

Gene-level count data from GSE171700 were used to examine ApoE-dependent transcription in isogenic human iPSC-derived neuron–astrocyte co-cultures. The primary analysis included four independent APOE ε3/ε3 and four APOE ε4/ε4 co-culture samples. Four APOE-knockout co-cultures were retained for secondary comparisons. Ensembl identifiers were stripped of version suffixes, mapped to human gene symbols using org.Hs.eg.db, and counts mapping to the same gene symbol were summed.

Genes with at least 10 reads in at least two samples were retained. Differential expression between APOE ε4/ε4 and ε3/ε3 co-cultures was analyzed using DESeq2 with genotype as the design factor. APOE ε3/ε3 was used as the reference, and log₂ fold changes were shrunk using the normal-prior estimator. Results were displayed as ε3/ε3 minus ε4/ε4 so that positive values indicate higher expression in APOE ε3/ε3. APOE-knockout comparisons were performed in a separate three-genotype model.

Variance-stabilized expression was calculated with blind=FALSE. For each predefined gene panel, expression was standardized within each gene across samples and averaged to generate one module score per independently cultured sample. Module scores were analyzed using linear models with genotype as the predictor. Gene-level log₂ fold-change distributions within each panel were tested against zero using two-sided Wilcoxon signed-rank tests. Benjamini-Hochberg correction was applied across the panel tests included in the external-validation analysis.

### Reanalysis of APOE4-to-APOE2 allelic-switch single-cell RNA sequencing

Processed 10x count matrices from GSE300079 were analyzed independently. The deposited experiment comprised three female mice with global APOE4-to-APOE2 switching, three with astrocyte-specific switching, one unswitched APOE4s2 control, two constitutive ApoE2-targeted-replacement mice and one constitutive ApoE4-targeted-replacement mouse. All animals received tamoxifen. Global-switch mice were 6-7 months old, astrocyte-switch mice were 4-5 months old, and the remaining deposited mice were 4 months old.

A Seurat object was generated independently for each mouse using genes detected in at least three cells. Sample-specific quality-control thresholds were calculated from median absolute deviations of log-transformed feature and count distributions. Cells were retained within three median absolute deviations below and four median absolute deviations above the sample medians, subject to a minimum of 200 detected genes and counts, a maximum of 7,500 genes, and a mitochondrial-read threshold of no more than 20%. Doublets were identified independently within each mouse using scDblFinder, and only singlets were retained.

Filtered samples were log-normalized and the 3,000 most variable genes were identified. Expression was scaled while regressing mitochondrial-read percentage, followed by principal-component analysis. Samples were integrated using reciprocal principal-component analysis with 3,000 integration features and dimensions 1–30. Nearest-neighbor graphs and UMAP coordinates were constructed from the first 30 integrated principal components, and clusters were identified at resolution 0.8. Cluster identities were assigned from standardized expression scores for canonical astrocyte, microglial, oligodendrocyte, oligodendrocyte precursor cell, endothelial, perivascular and neuronal markers and were reviewed using marker dot plots and cluster -enriched genes.

Differential-expression testing used unintegrated raw RNA counts. Counts were summed within each mouse and cell population to generate mouse-level pseudobulk profiles. Astrocytes, microglia, oligodendrocytes, oligodendrocyte precursor cells, endothelial cells and neurons were analyzed. Cluster 38 was excluded from the primary oligodendrocyte population, and cluster 27 was excluded from the primary endothelial population because of mixed endothelial and pericyte/vascular smooth-muscle marker expression; an endothelial analysis retaining cluster 27 was performed as a sensitivity analysis. Populations required at least 100 cells from each mouse, except oligodendrocyte precursor cells, for which the threshold was 30 cells.

Pseudobulk counts were analyzed using edgeR. Genes were filtered using filterByExpr with a minimum count of five, normalized by the trimmed mean of M-values method, and fitted using robust quasi-likelihood negative-binomial models containing all five experimental groups. Prespecified contrasts compared global-switch with unswitched APOE4s2, astrocyte-switch with unswitched APOE4s2, and constitutive ApoE2-targeted-replacement with ApoE4-targeted-replacement mice. Positive coefficients indicate higher expression in the first group named by each contrast.

Coordinated differential expression of the predefined antioxidant, inflammatory, pro-oxidant and external Nrf2 target sets was tested using CAMERA, with inter-gene correlation incorporated into the competitive gene-set test. Benjamini-Hochberg correction was applied across all cell-type, contrast and panel tests. Sample-level module scores were calculated as the mean of gene-wise standardized TMM-normalized log₂ counts per million. Gene-level Wilcoxon and sign tests were retained as descriptive analyses and were not used as substitutes for mouse-level inference.

### Reanalysis of cognitively normal human single-nucleus RNA sequencing

Processed single-nucleus count matrices from GSE237718 were reanalyzed using cognitively normal donors only. The analysis included 27 donors: 9 APOE ε2/ε3, 10 ε3/ε3, 5 ε3/ε4 and 3 ε4/ε4 individuals. Each donor was retained as the biological unit.

Seurat objects were generated separately for each donor. Nuclei with 200-10,000 detected genes and no more than 5% mitochondrial reads were retained, and doublets were identified using scDblFinder. Singlet nuclei were merged, log-normalized using a scale factor of 10,000, and the 3,000 most variable genes were identified. Expression was scaled while regressing detected-feature number and mitochondrial-read percentage, followed by principal-component analysis. Harmony^64^ correction by donor was applied to the first 30 principal components for clustering and visualization only. Clusters were identified at resolution 0.4 and annotated as astrocytes, excitatory neurons, inhibitory neurons, microglia, oligodendrocytes, oligodendrocyte precursor cells, endothelial cells or pericytes using canonical marker expression.

Raw counts were summed by donor within each cell type. A donor-cell-type profile required at least 30 nuclei for astrocytes, neuronal populations, microglia, oligodendrocytes and oligodendrocyte precursor cells, 20 nuclei for endothelial cells, or 15 nuclei for pericytes. Cell types represented by fewer than eight donors were not tested.

Pseudobulk counts were analyzed using edgeR^65^. Genes were retained using filterByExpr with a minimum count of 10, normalized by the trimmed mean of M-values method, and fitted using robust quasi-likelihood models. Sex was included as a covariate when the resulting design matrix remained full rank. Th e primary exact-genotype model compared APOE ε2/ε3, ε3/ε4 and ε4/ε4 separately with ε3/ε3. A complementary three-group model combined ε3/ε4 and ε4/ε4 donors as APOE ε4 carriers. An ordered-genotype model coded ε4/ε4 as -2, ε3/ε4 as -1, ε3/ε3 as 0 and ε2/ε3 as +1. Competitive panel enrichment was tested with CAMERA using signed quasi-likelihood statistics, calculated as the sign of the log_2_ fold change multiplied by the square root of the F statistic. Benjamini-Hochberg correction was applied within each contrast across all evaluable cell-type-by-panel tests. Donor-level module scores were calculated by gene-wise standardization of TMM-normalized log_2_ counts per million with a prior count of two, followed by averaging across represented genes.

Genes contributing recurrently to the antioxidant-panel result were identified descriptively by counting the cell types in which each gene was lower in APOE ε4 carriers, calculating its median log_2_ fold change and mean signed statistic across cell types, and examining directional concordance in the separate ε3/ε4 and ε4/ε4 contrasts. This cross-cell-type ranking was used to identify contributors to the panel-level effect and was not treated as gene-level differential-expression significance.

### AMP-AD Diverse Cohorts bulk RNA sequencing

Gene-level counts and harmonized metadata were obtained from the AMP-AD Diverse Cohorts Study through the AD Knowledge Portal, accession syn68755624^31^. This resource integrates data contributed by Columbia, Emory, Mayo, Mount Sinai and Rush. Analyses were restricted to neuropathologically defined controls carrying APOE ε2/ε3, ε3/ε3 or ε3/ε4. Control selection used harmonized CERAD neuritic-plaque classifications and Braak staging, incorporating Mayo diagnostic classifications as specified in the resource. Eligible categories included absent or sparse/infrequent neuritic plaques, including C0 or C1 classifications, with Braak stage ≤III.

Caudate nucleus, dorsolateral prefrontal cortex and superior temporal gyrus were analyzed separately. Final ε2/ε3, ε3/ε3 and ε3/ε4 specimen counts were 9, 56 and 7 for caudate; 15, 93 and 16 for dorsolateral prefrontal cortex; and 20, 112 and 17 for superior temporal gyrus.

Ensembl identifiers were mapped to gene symbols using GENCODE v43/GRCh38 annotations, and counts were rounded to integers. Genes required at least 10 counts in at least 10 samples within each region. Differential expression used DESeq2 1.38.3 in R 4.2.3, with median-of-ratios normalization. Models included APOE genotype, a natural spline of standardized RIN with two degrees of freedom, PMI group, sex, race, Hispanic ethnicity and data-contribution group. PMI was categorized as ≤5 hours, >5–10 hours, >10 hours or missing.

Wald tests evaluated ε2/ε3 versus ε3/ε3, ε3/ε4 versus ε3/ε3 and ε2/ε3 versus ε3/ε4. A separate ordinal model replaced categorical genotype with ε3/ε4=−1, ε3/ε3=0 and ε2/ε3=+1 while retaining the same covariates. Positive coefficients indicate increasing expression toward ε2/ε3 and negative coefficients toward ε3/ε4. Benjamini-Hochberg correction was applied genome-wide within each region and contrast. Predefined panel genes were extracted from the complete results irrespective of significance.

### ROSMAP prefrontal-cortex single-nucleus RNA sequencing

Raw single-nucleus counts and donor metadata were obtained from the MIT ROSMAP Single-Nucleus Multiomics Study through the AD Knowledge Portal, accession syn52392369^32^. Inclusion required a final consensus clinical diagnosis of no cognitive impairment at death, defined as no impaired cognitive domains, APOE ε2/ε3, ε3/ε3 or ε3/ε4 genotype, and complete age and sex information^66^.

Raw counts were summed within each donor–major-cell-type combination. Profiles required at least 30 nuclei for astrocytes, excitatory neurons, inhibitory neurons, microglia, oligodendrocytes and oligodendrocyte precursor cells; 20 nuclei for endothelial cells; and 15 nuclei for pericytes, CNS-associated macrophages, fibroblasts or T cells. Smooth-muscle-cell profiles were excluded. Pseudobulk columns were aligned to metadata by donor identifier. The six principal populations included 141 astrocyte, 143 excitatory-neuron, 141 inhibitory-neuron, 133 microglial, 143 oligodendrocyte and 140 oligodendrocyte-precursor-cell donor profiles. Genotype-specific counts are provided with the source data.

Differential expression was performed separately by cell type using edgeR 3.40.2 in R 4.2.3. Genes were filtered using filterByExpr with min.count = 10. Counts were TMM-normalized, dispersions were estimated using robust empirical-Bayes estimation, and robust quasi-likelihood negative-binomial models were fitted using glmQLFit. Categorical models included age, sex and APOE genotype. Quasi-likelihood F-tests evaluated ε2/ε3 versus ε3/ε3, ε3/ε4 versus ε3/ε3 and ε2/ε3 versus ε3/ε4.

A separate ordinal model replaced categorical genotype with ε3/ε4 = −1, ε3/ε3 = 0 and ε2/ε3 = +1 while retaining age and sex. All expressed genes were included during model fitting and genome-wide Benjamini– Hochberg adjustment. Predefined panel genes were subsequently extracted without selecting on significance. Positive ordinal coefficients indicate an ε2/ε3 bias and negative coefficients an ε3/ε4 bias.

### Statistical analysis

Statistical analyses were performed in GraphPad Prism 11, SPSS version 29.0.2.0 and R 4.3.2. Exact sample sizes and statistical tests are reported in the figure legends and source data. Two independent groups were compared by two-tailed unpaired t-test or Mann-Whitney test as appropriate. More than two independent groups were analyzed by one-way ANOVA with Tukey correction or Kruskal-Wallis test with Dunn correction. Experiments with two factors were analyzed by two-way ANOVA with Tukey correction. Matched gene-level distributions were analyzed by Friedman tests with Dunn correction or paired Wilcoxon procedures as indicated. Coefficient distributions were tested against zero by two-sided Wilcoxon signed-rank tests. Positional distributions used Stuart-Maxwell tests, and continuous associations used two-sided Spearman correlation. Module differences were assessed by permutation; gene-set overlaps used two-sided Fisher exact tests. Benjamini-Hochberg correction was used unless otherwise specified. All tests were two-sided, and *P*<0.05 or adjusted *P*<0.05 was considered significant. Data are shown as individual measurements with mean±SEM; gene-level distributions are summarized by median and interquartile range.

### Data availability

The RNA-sequencing datasets generated in this study are available through ArrayExpress under E-MTAB-16242 (ApoE-isoform U87 cells treated with DMSO or SFN) and E-MTAB-14167 (human prefrontal cortex and cerebellum). Public datasets reanalyzed in this study are available through the AD Knowledge Portal/Synapse (aging ApoE-targeted-replacement mouse cortex, syn15682620) and NCBI Gene Expression Omnibus (isogenic human iPSC-derived neuron-astrocyte co-cultures, GSE171700; APOE4-to-APOE2 allelic-switch single-cell RNA sequencing, GSE300079; and cognitively normal human temporal-cortex single-nucleus RNA sequencing, GSE237718). AMP-AD Diverse Cohorts bulk RNA-sequencing data and MIT ROSMAP prefrontal-cortex single-nucleus RNA-sequencing data were accessed through the AD Knowledge Portal/Synapse under syn68755624 and syn52392369, respectively. Access to controlled human data is subject to the requirements of the AD Knowledge Portal and contributing studies. The Tabula Muris Senis reference used for MuSiC deconvolution is available through GEO under GSE149590, within SuperSeries GSE132042. The published Nrf2 target sets used for regulon analysis were obtained from Malhotra et al.^51^ The authors declare that the data supporting the findings of this study are available within the paper and its Supplementary Information files. Should any raw data files be needed in another format, they are available from the corresponding author upon reasonable request.

---

**Extended Data Figure 1.**
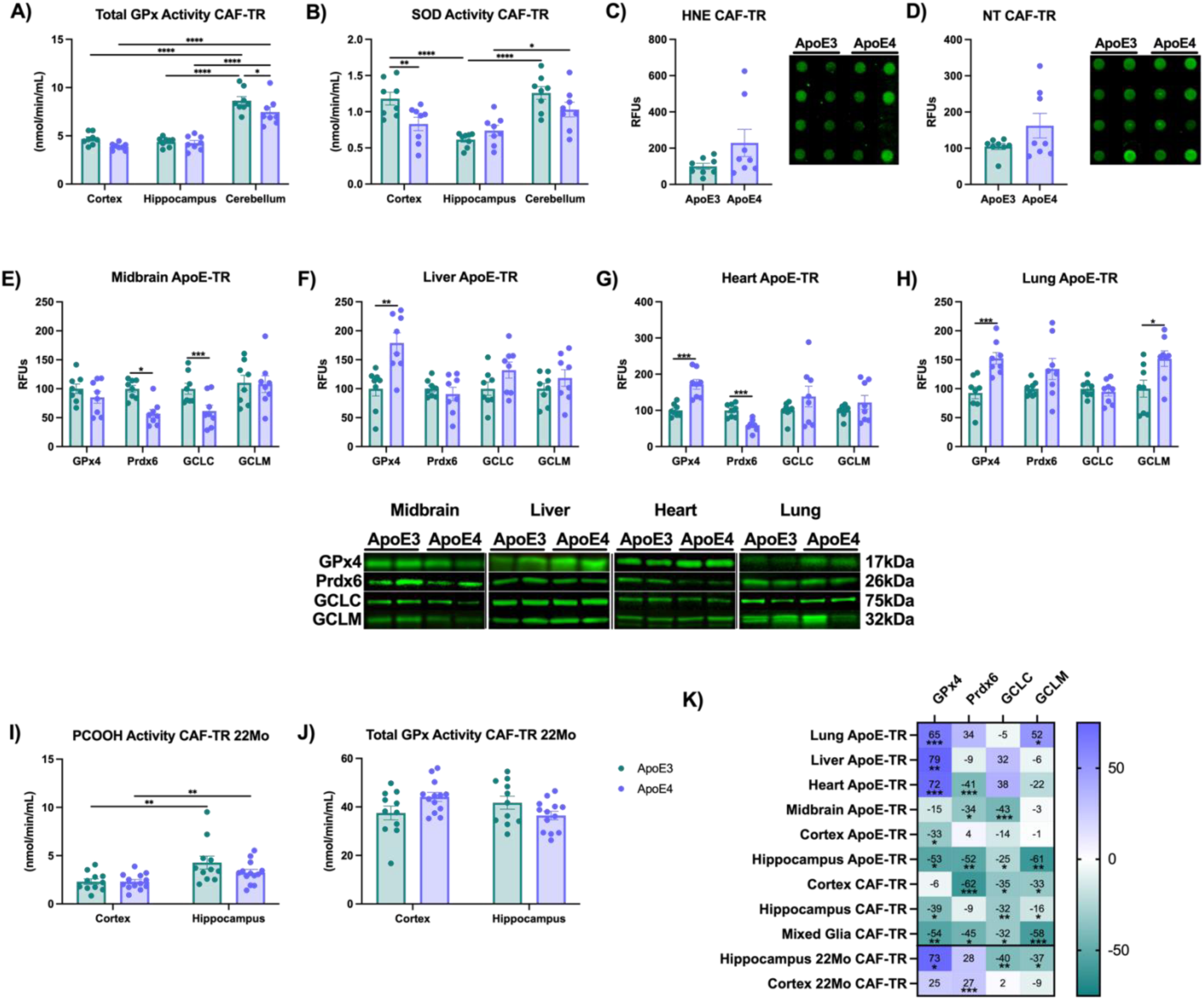
Regional, peripheral and age-dependent antioxidant phenotypes across humanized ApoE targeted-replacement mouse models. **A)** Total glutathione peroxidase (GPx) activity and **B)** superoxide dismutase (SOD) activity in cortex, hippocampus and cerebellum from 6-month-old CAF-TR mice (n=4/sex/group). Oxidative damage markers **C)** HNE and **D)** nitrotyrosine (NT) in 6-month-old CAF-TR brain tissue. GPx4, Prdx6, GCLC and GCLM protein abundance in **E)** midbrain, **F)** liver, **G)** heart and **H)** lung from 6-month-old ApoE-TR mice. **I)** Oxidized phosphatidylcholine (PCOOH) reducing activity and **J)** total GPx activity in cortex and hippocampus from 22-month-old CAF-TR mice (n=4-8/sex/group). **K)** Summary heatmap of ApoE4-associated differences in GPx4, Prdx6, GCLC and GCLM protein abundance across brain regions, peripheral tissues, primary mixed glia, mouse models and ages; values are percent difference in ApoE4 relative to ApoE3, with positive values indicating higher and negative values lower abundance in ApoE4. Within each mouse model, tissues and brain regions were collected from the same animals used in Figure 1. Protein abundance, HNE and NT are expressed as relative fluorescence units (RFU). Data are shown as mean±SEM. A,B,I,J) by two-way ANOVA with ApoE isoform and brain region as factors followed by Tukey’s multiple-comparisons test. C-H) by two-tailed unpaired t-test or Mann-Whitney test as appropriate. *p<0.05, **p<0.01, ***p<0.001, ****p<0.0001.

**Extended Data Figure 2.**
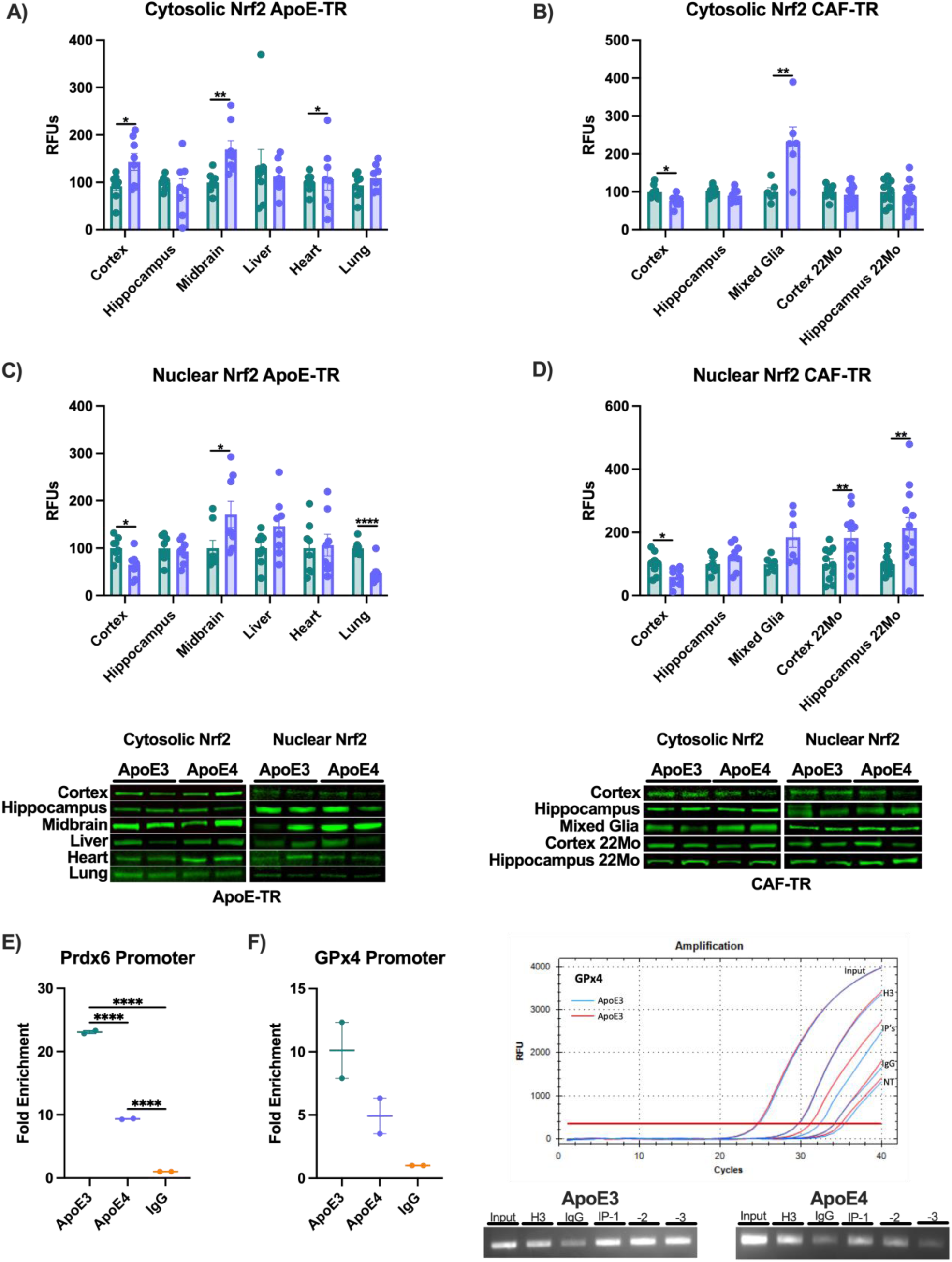
ApoE-dependent Nrf2 localization and antioxidant-promoter occupancy across tissues and aging. Cytosolic Nrf2 abundance in **A)** tissues from 6-month-old ApoE-TR mice and **B)** CAF-TR brain regions, primary mixed glia and aged brain. Corresponding nuclear Nrf2 abundance in **C)** ApoE-TR and **D)** CAF-TR samples. The CAF-TR panels include 6-month-old cortex and hippocampus, neonatal primary cortical mixed glia, and 22-month-old cortex and hippocampus. Nrf2 occupancy at the **E)** Prdx6 and **F)** GPx4 promoter regions in cortex from 6-month-old CAF-TR mice, measured by ChIP-qPCR and expressed relative to rabbit IgG. Three cortices were pooled per biological replicate, with two independent pools per genotype. Representative immunoblots, amplification curves and promoter-PCR products are shown. Protein and ChIP-qPCR data are presented as individual biological measurements with mean±SEM. A-D) Two-tailed unpaired t-tests or Mann-Whitney tests, as appropriate. E,F) One-way ANOVA with Tukey’s multiple-comparisons test. *P<0.05, **P<0.01, ***P<0.001, ****P<0.0001.

**Extended Data Figure 3.**
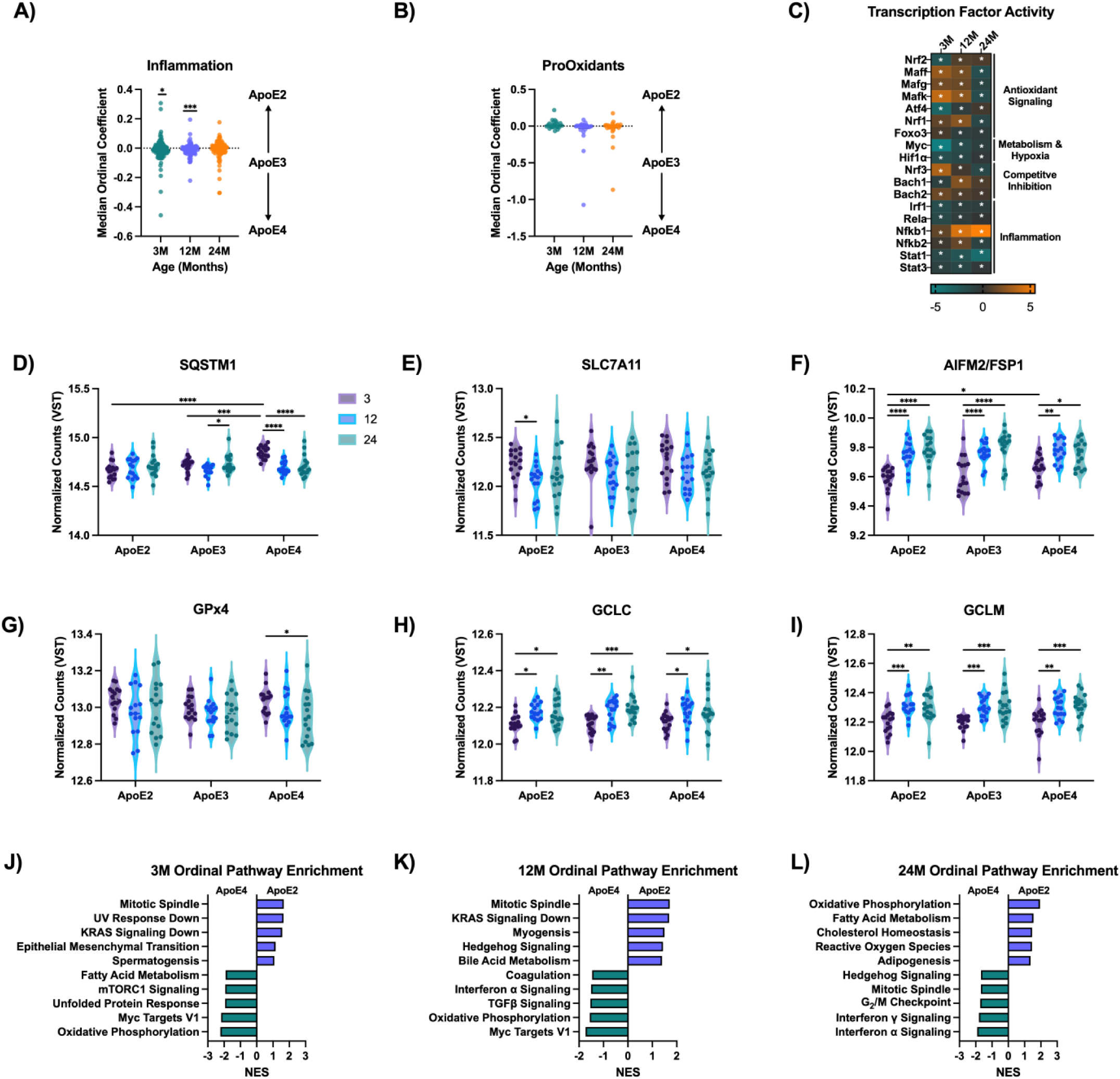
Age-specific transcriptional responses along the ApoE ordinal axis. Cortical RNA-sequencing data from the ApoE-targeted-replacement aging cohort were analyzed using ApoE4→ApoE3→ApoE2 ordinal coding; positive coefficients indicate ApoE2-biased expression and negative coefficients ApoE4-biased expression. Ordinal-coefficient distributions for the predefined **A)** inflammatory and **B)** pro-oxidant panels at 3, 12 and 24 months. **C)** DoRothEA-inferred activity of transcription factors associated with antioxidant regulation, metabolism, hypoxia and inflammatory signaling at each age. VST-normalized expression of **D)** SQSTM1, **E)** SLC7A11, **F)** AIFM2/FSP1, **G)** GPx4, **H)** GCLC and **I)** GCLM across ApoE isoforms and ages. Transcriptome-wide Hallmark GSEA at **J)** 3, **K)** 12 and **L)** 24 months, displayed as normalized enrichment scores. In A,B, each point represents one gene, with median and interquartile range; D-I show sample-level expression distributions. A,B) Two-sided Wilcoxon signed-rank tests against zero and Friedman tests with Dunn’s multiple-comparisons tests across ages. C) Permutation tests with Benjamini-Hochberg correction. D-I) Two-way ANOVA with age, ApoE isoform and their interaction, followed by Tukey’s multiple-comparisons tests. J-L) GSEA with false discovery rate correction. *P<0.05, **P<0.01, ***P<0.001, ****P<0.0001.

**Extended Data Figure 4.**
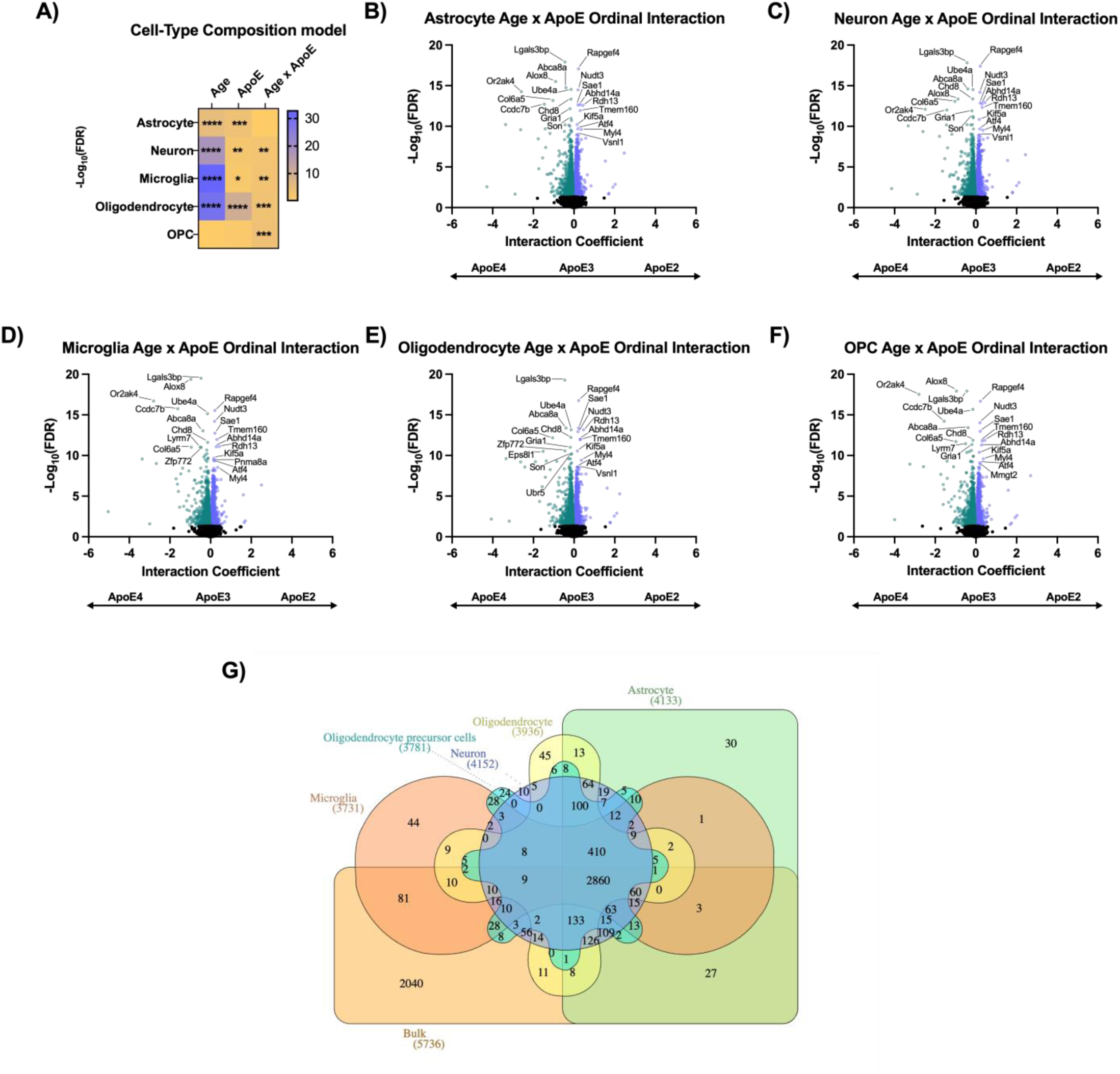
Cell-type-enriched Age x ApoE ordinal interactions and cross-analysis overlap. **A)** Heatmap of −log_10_(FDR) for age, ApoE genotype and Age x ApoE effects on MuSiC-estimated cell-type proportions. Endothelial and pericyte estimates were excluded from downstream adjustment because their estimated proportions were negligible. MuSiC-derived cell-type-enriched Age x ApoE ordinal interaction volcano plots for **B)** astrocytes, **C)** neurons, **D)** microglia, **E)** oligodendrocytes and **F)** oligodendrocyte precursor cells (OPCs); each point represents a gene, and the ordinal scale beneath each panel indicates the direction ApoE4→ApoE3→ApoE2. **G)** Venn diagram of overlap among the bulk and five cell-type-enriched Age x ApoE ordinal interaction DEG sets. A) by linear models containing age, genotype, age x genotype and sex, with false discovery rate correction. B-F) by DESeq2 Wald test with Benjamini-Hochberg correction.

**Extended Data Figure 5.**
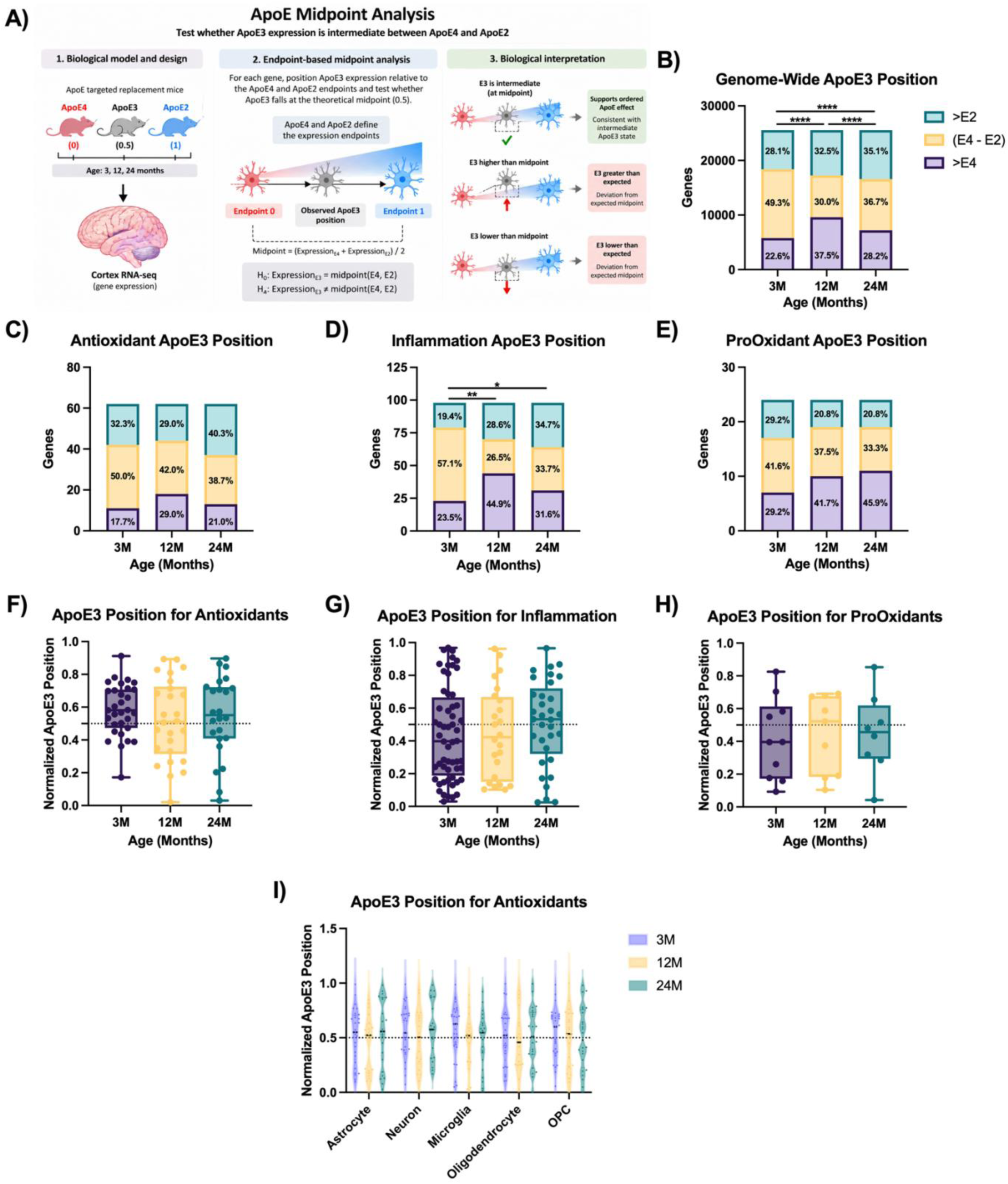
ApoE3 midpoint analysis defines age- and pathway-dependent positioning between ApoE4 and ApoE2 transcriptional states. **A)** Schematic of the ApoE3 midpoint analysis. ApoE isoforms were modeled categorically at each age, with ApoE4 and ApoE2 defining the expression endpoints and ApoE3 position calculated relative to these endpoints; a normalized position of 0 corresponds to Apo E4, 1 to ApoE2 and 0.5 to the theoretical midpoint. **B)** Genome-wide classification of ApoE3 position at 3, 12 and 24 months as beyond ApoE4, between ApoE4 and ApoE2, or beyond ApoE2. Corresponding positional-class distributions for the predefined **C)** antioxidant, **D)** inflammatory and **E)** pro-oxidant gene sets. Normalized ApoE3 position among genes classified as intermediate between ApoE4 and ApoE2 for the **F)** antioxidant, **G)** inflammatory and **H)** pro-oxidant modules. **I)** Normalized ApoE3 positions for intermediate antioxidant-panel genes in astrocyte-, neuron-, microglia-, oligodendrocyte- and OPC-enriched profiles at 3, 12 and 24 months. In F-I, the dotted line denotes the theoretical midpoint (0.5). Distribution plots show individual genes with median and interquartile range. B) by paired Stuart-Maxwell test with Holm correction for pairwise age comparisons. C-E) by paired Stuart-Maxwell test with Benjamini-Hochberg correction across module comparisons; antioxidant and pro-oxidant distributions did not redistribute with age, whereas inflammatory distributions differed between 3 and 12 months and between 3 and 24 months. F-I) by two-sided Wilcoxon signed-rank test against 0.5 with Benjamini-Hochberg correction; no distribution differed from the midpoint after correction.

**Extended Data Figure 6.**
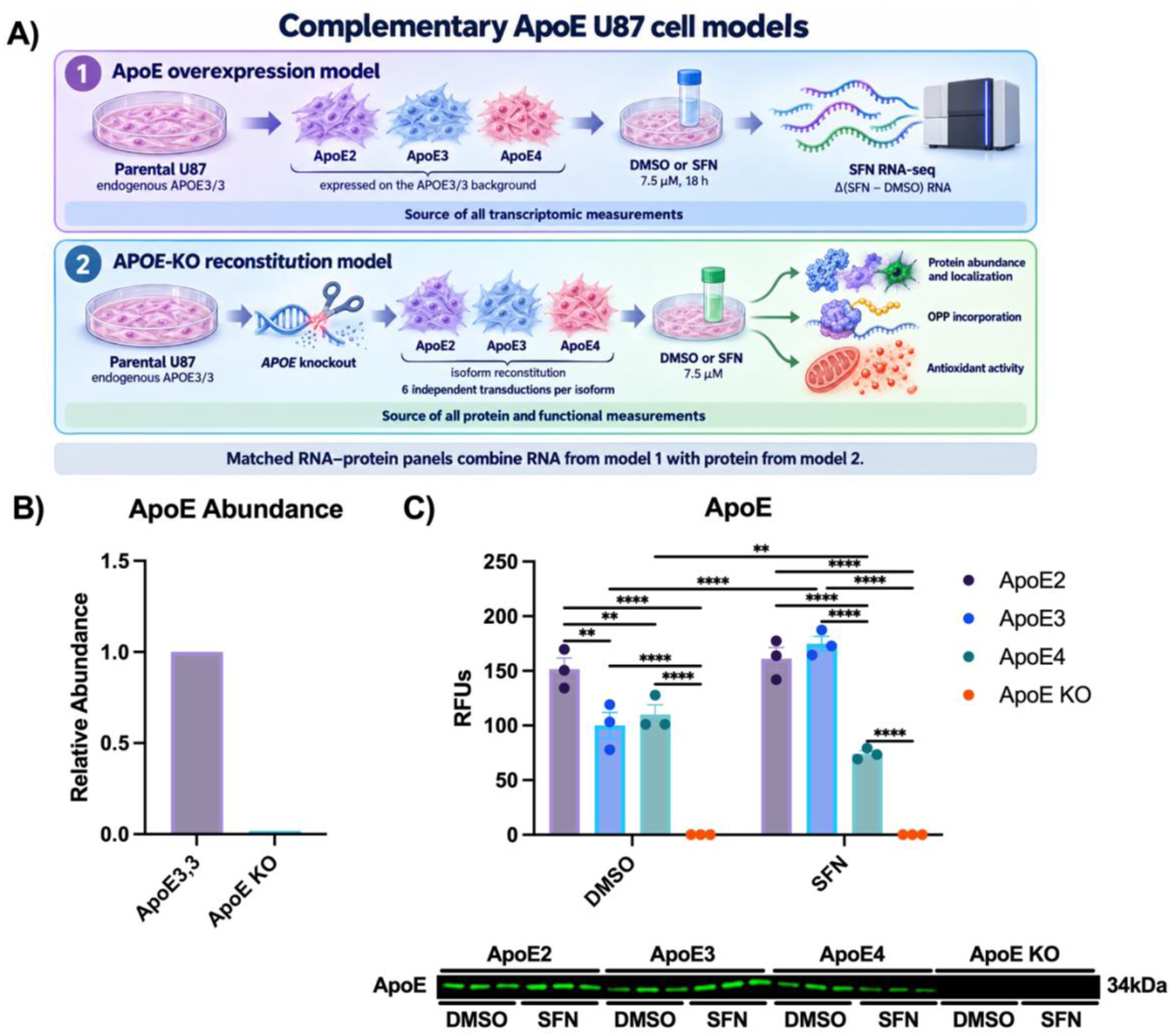
Generation and validation of complementary ApoE-expressing U87 cell models. **A)** Experimental design for the two independently generated U87 models. In the first model, APOE2, APOE3 or APOE4 was expressed on the endogenous APOE3/3 background and used for SFN RNA sequencing and all Δ(SFN−DMSO) RNA measurements. In the second model, endogenous APOE was disrupted and APOE-KO cells were independently reconstituted with APOE2, APOE3 or APOE4 using six independent transductions per isoform; this model was used for all protein, OPP-incorporation and antioxidant-activity measurements. Matched RNA-protein panels therefore combine RNA measurements from the first model with protein measurements from the second. **B)** Relative ApoE abundance detected by mass spectrometry in parental ApoE3 and ApoE KO U87 cells, normalized to ApoE3. **C)** ApoE abundance in RIPA lysates of ApoE2, ApoE3, ApoE4 and ApoE KO cells following DMSO or 7.5μM SFN for 18 hours; a representative immunoblot is shown below. Data are shown as individual biological replicates with mean±SEM. C) by two-way ANOVA with ApoE background and treatment as factors followed by Tukey’s multiple-comparisons test. *p<0.05, **p<0.01, ***p<0.001, ****p<0.0001.

**Extended Data Figure 7.**
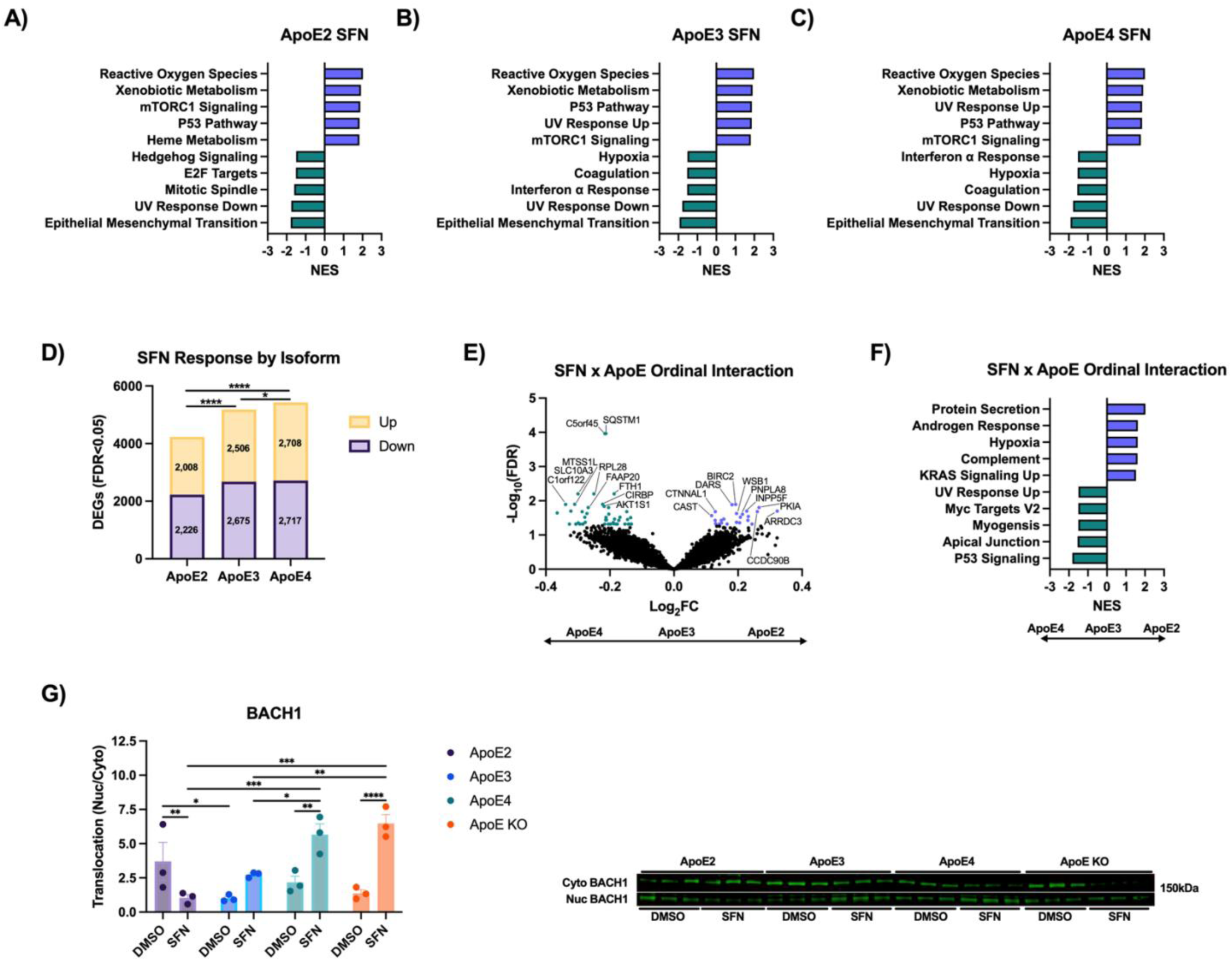
Extended transcriptomic and regulatory analyses of the ApoE-dependent SFN response. GSEA of SFN-induced transcriptional changes in **A)** ApoE2, **B)** ApoE3 and **C)** ApoE4 cells, showing the top positively and negatively enriched Hallmark pathways as normalized enrichment scores (NES). **D)** Genome-wide breadth of the SFN response across ApoE isoforms; bars show the numbers of upregulated and downregulated DEGs (ApoE2, 4,234; ApoE3, 5,181; ApoE4, 5,425). **E)** Volcano plot of the SFN x ApoE ordinal interaction, with isoforms ordered ApoE4→ApoE3→ApoE2. **F)** GSEA of the ranked SFN x ApoE ordinal interaction statistic, showing the top positively and negatively enriched pathways. **G)** BACH1 translocation following DMSO or SFN treatment in ApoE2, ApoE3, ApoE4 and ApoE KO cells; representative immunoblots are shown below. Protein data are shown as mean±SEM with individual values overlaid. A-C,F) by GSEA with false discovery rate correction. D) gene-state distributions by paired Stuart-Maxwell test with Holm correction (ApoE2 vs ApoE3, adjusted *P*=2.58x10^−21^; ApoE2 vs ApoE4, adjusted *P*=1.79x10^−33^; ApoE3 vs ApoE4, adjusted *P*=0.0169). E) by DESeq2 Wald test with Benjamini-Hochberg correction. G) by two-way ANOVA with ApoE genotype and treatment as factors followed by Tukey’s multiple-comparisons test. *p<0.05, **p<0.01, ***p<0.001, ****p<0.0001.

**Extended Data Figure 8.**
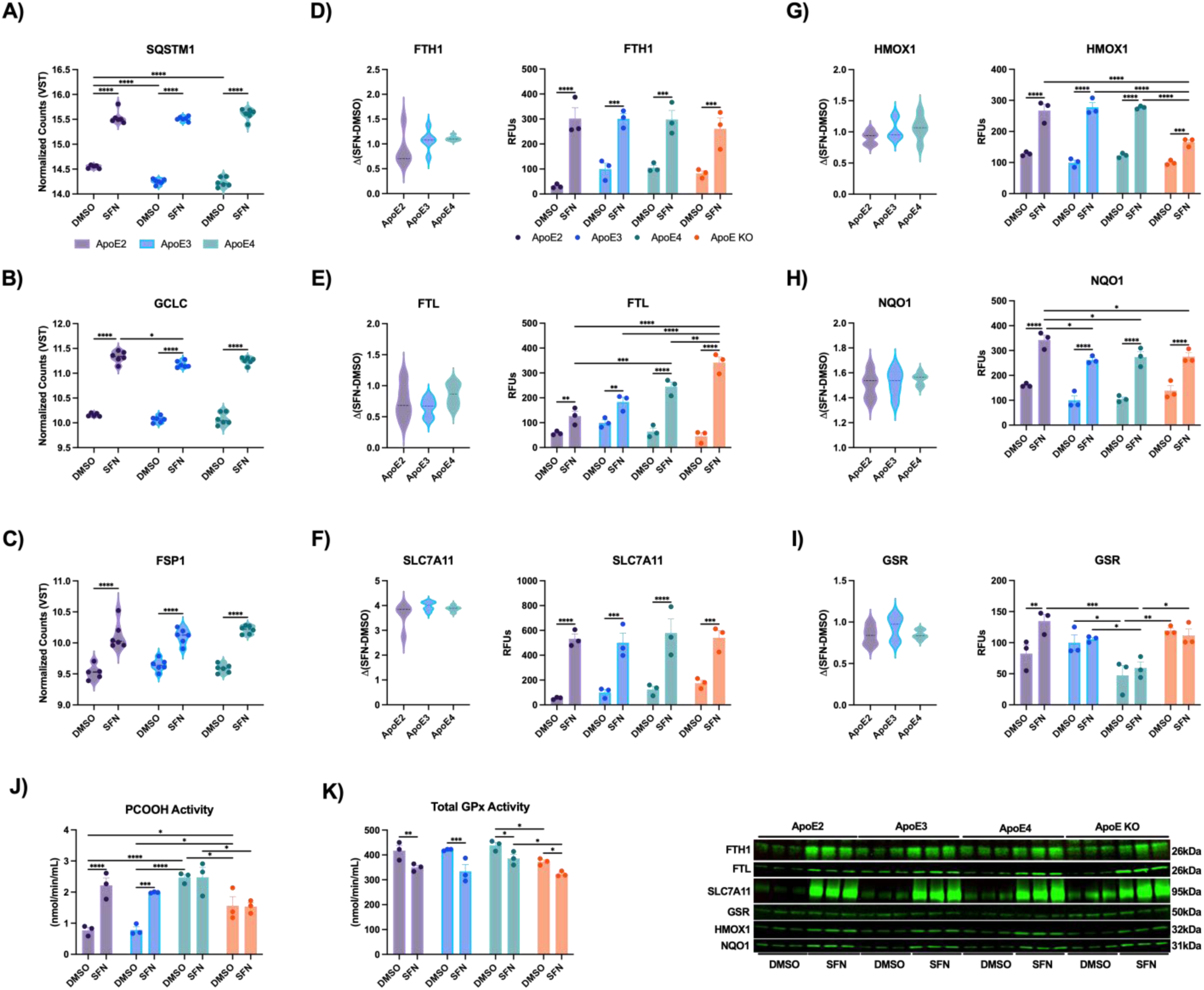
Expanded analysis of ApoE-dependent sulforaphane responses across antioxidant transcripts, proteins and antioxidant activity. VST-normalized RNA expression of **A)** SQSTM1, **B)** GCLC and **C)** FSP1 following DMSO or SFN treatment. Matched RNA-protein analyses for **D)** FTH1, **E)** FTL, **F)** SLC7A11/xCT, **G)** HMOX1, **H)** NQO1 and **I)** GSR; for each gene, the left panel shows the SFN-induced RNA change [Δ(SFN−DMSO)] and the right panel the corresponding protein abundance following DMSO or SFN treatment. **J)** PCOOH-reducing and **K)** total GPx activity following DMSO or SFN treatment. Representative immunoblots are shown below the quantitative panels. Data are shown as individual measurements with mean±SEM. A-C) by two-way ANOVA with ApoE genotype and treatment as factors followed by Tukey’s multiple-comparisons test. D-I) ΔRNA by one-way ANOVA with Tukey’s multiple-comparisons test; protein by two-way ANOVA with ApoE genotype and treatment as factors followed by Tukey’s multiple-comparisons test. J,K) by two-way ANOVA with ApoE genotype and treatment as factors followed by Tukey’s multiple-comparisons test. *p<0.05, **p<0.01, ***p<0.001, ****p<0.0001.

**Extended Data Figure 9.**
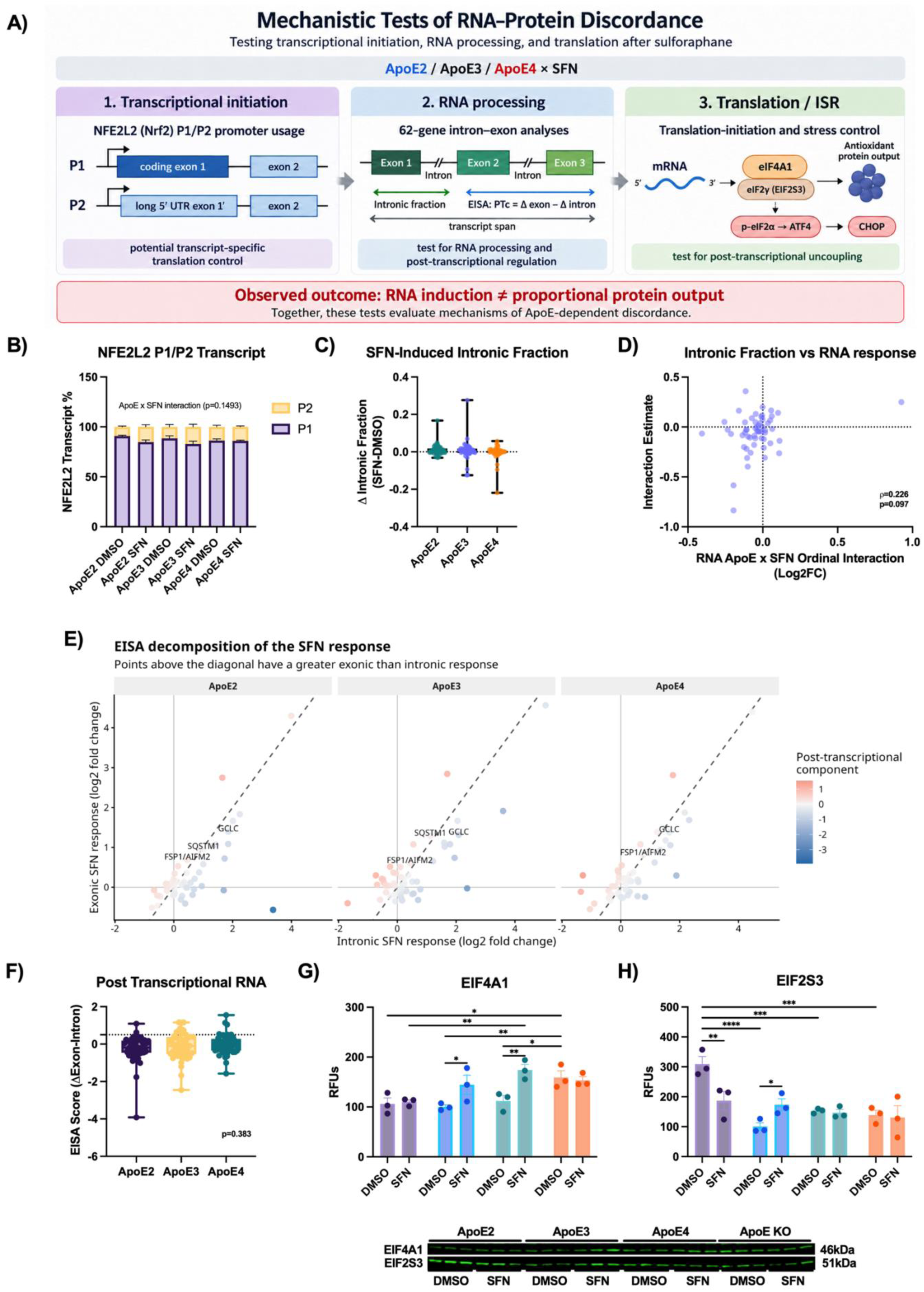
Mechanistic tests of ApoE-dependent RNA–protein discordance. U87 cells expressing ApoE2, ApoE3 or ApoE4 were treated with DMSO or 7.5 μM SFN for 18 hours. **A)** Schematic of the mechanisms examined: NFE2L2 promoter usage, intronic fraction, exon–intron split analysis and translation-initiation machinery. **B)** Relative contributions of P1- and P2-derived NFE2L2 transcripts estimated from length-normalized reads assigned to promoter-specific first exons. **C)** SFN-induced change in intronic fraction, calculated as intronic/(intronic+exonic) reads, across the antioxidant-panel genes. **D)** Relationship between RNA SFN x ApoE ordinal interaction coefficients and corresponding intronic-fraction interaction estimates. **E)** Exon-intron split analysis comparing exonic and intronic SFN responses across antioxidant genes; each point represents one gene, the dashed diagonal denotes equivalent responses, and color indicates the post- transcriptional component. **F)** Corresponding post-transcriptional component, calculated as the exonic SFN response minus the intronic response. Protein abundance of **G)** EIF4A1 and **H)** EIF2S3 following DMSO or SFN treatment in ApoE2, ApoE3, ApoE4 and ApoE KO cells. Representative immunoblots are shown below. Gene-level distributions show individual genes with median and interquartile range; protein measurements show individual biological replicates with mean±SEM. B) Quasibinomial SFN x ApoE ordinal interaction model. C) Friedman test across isoforms. D) Two-sided Spearman correlation. F) Friedman test across isoforms and two-sided Wilcoxon signed-rank tests against zero with Benjamini-Hochberg correction. G,H) Two-way ANOVA with ApoE background and treatment as factors, followed by Tukey’s multiple-comparisons tests. *P<0.05, **P<0.01, ***P<0.001, ****P<0.0001.

**Extended Data Figure 10.**
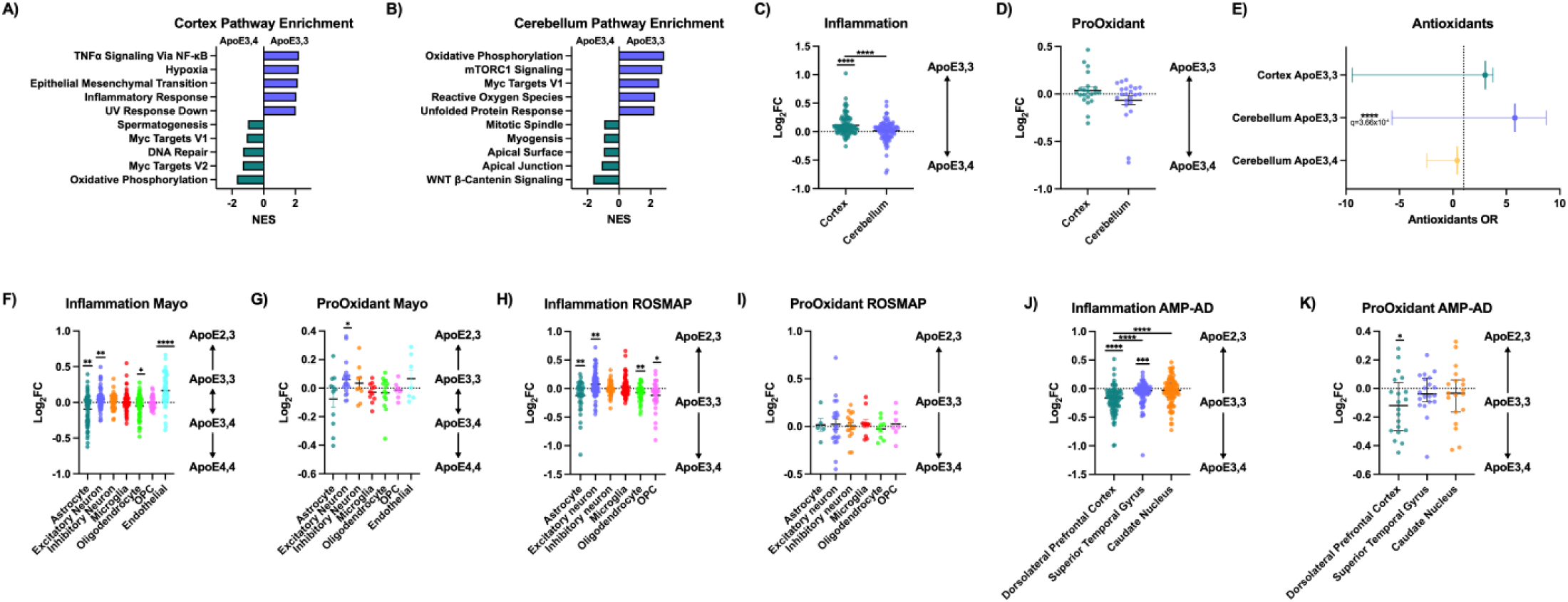
APOE-associated pathway and gene-panel relationships across human brain regions and cell types. Hallmark GSEA in **A)** cortex and **B)** cerebellum from cognitively normal APOE ε3/ε3 and ε3/ε4 adults. Positive normalized enrichment scores indicate an ε3/ε3 bias and negative scores an ε3/ε4 bias. Regional log_2_ fold-change distributions for the predefined **C)** inflammatory and **D)** pro-oxidant panels; positive values indicate higher expression in ε3/ε3. **E)** Antioxidant-panel enrichment among directional FDR-significant transcripts, shown as odds ratios with 95% confidence intervals. The cortical ε3/ε4-biased set contained no antioxidant-panel genes and was omitted. Gene-level ordinal coefficients for the **F)** inflammatory and **G)** pro-oxidant panels across temporal-cortex cell populations from 27 donors without AD neuropathology from the Mayo Clinic Brain Bank cohort. Genotypes were ordered ε4/ε4→ε3/ε4→ε3/ε3→ε2/ε3; positive coefficients indicate an ε2/ε3 bias and negative coefficients an ε4/ε4 bias. Corresponding **H)** inflammatory and **I)** pro-oxidant coefficients across ROSMAP prefrontal-cortex cell populations from clinically cognitively unimpaired donors, and **J)** inflammatory and **K)** pro-oxidant coefficients across AMP-AD brain regions from donors without AD neuropathology. Distribution plots show individual genes with median and interquartile range; dotted horizontal lines denote zero. A,B) GSEA with false discovery rate correction. C,D) Two-sided Wilcoxon signed-rank tests against zero within regions and Wilcoxon matched-pairs signed-rank tests between regions. E) Two-sided Fisher’s exact tests with Benjamini-Hochberg correction. F-K) Two-sided Wilcoxon signed-rank tests against zero; regional distributions in J) were additionally compared using a Friedman test with Dunn’s multiple-comparisons test. *P<0.05, **P<0.01, ***P<0.001, ****P<0.0001.

**Supplementary Table 1.** Characteristics of cognitively normal human brain samples used for RNA-sequencing and protein analyses. Demographic and neuropathological characteristics are shown for prefrontal cortex and cerebellum samples from individuals with *APOE* ε3/ε3 or ε3/ε4 genotypes. Each row represents an individual tissue specimen. Samples marked “Yes” in the RNA-seq column were included in the transcriptomic analysis; samples marked “No” were used for western blotting. “Both regions available” indicates that prefrontal cortex and cerebellum were available from the same donor. RIN is reported only for samples used for RNA-seq. PMI, postmortem interval; RIN, RNA integrity number; MMSE, Mini-Mental State Examination.

| Brain region | *APOE* genotype | Sex | Age | Braak | PMI (hours) | MMSE | MMSE interval (months) | Bank | RNA-seq | RIN | Both regions available |
|---|---|---|---|---|---|---|---|---|---|---|---|
| Prefrontal cortex | ε3/ε3 | Female | 99 | 0 | 9 | 28 | 23.4 | USC | No | — | Yes |
| Prefrontal cortex | ε3/ε3 | Female | 91 | 0 | 8.75 | 20 | 84 | USC | Yes | 4.6 | Yes |
| Prefrontal cortex | ε3/ε3 | Female | 85 | 0 | 7 | — | — | USC | Yes | 4.2 | Yes |
| Prefrontal cortex | ε3/ε3 | Female | 95 | 1 | 3.25 | — | — | USC | Yes | 6 | Yes |
| Prefrontal cortex | ε3/ε3 | Female | 87 | 2 | 10.75 | 29 | 1 | USC | Yes | 2.9 | Yes |
| Prefrontal cortex | ε3/ε3 | Female | 95 | 2 | 2.92 | — | — | UCI | Yes | 5.2 | Yes |
| Prefrontal cortex | ε3/ε3 | Female | 86 | 3 | 6.17 | 30 | 35.5 | UCI | Yes | 2.4 | Yes |
| Prefrontal cortex | ε3/ε3 | Female | 90 | 3 | 2.2 | — | — | UCI | Yes | 5.9 | Yes |
| Prefrontal cortex | ε3/ε3 | Female | 89 | 3 | 3.58 | 24 | 5.9 | UCI | Yes | 4.5 | No |
| Prefrontal cortex | ε3/ε4 | Female | 91 | 3 | 11.2 | 23 | 10.1 | UW | No | — | Yes |
| Prefrontal cortex | ε3/ε4 | Female | 91 | 3 | 3.33 | 27 | 42.5 | UCI | No | — | Yes |
| Prefrontal cortex | ε3/ε4 | Female | 87 | 3 | 4.57 | 29 | 85.6 | UCI | Yes | 2.6 | Yes |
| Prefrontal cortex | ε3/ε4 | Female | 93 | 5 | 4.15 | 30 | 81.6 | UCI | Yes | 4 | Yes |
| Prefrontal cortex | ε3/ε3 | Male | 82 | 0 | 9 | 28 | — | USC | Yes | 6.2 | Yes |
| Prefrontal cortex | ε3/ε3 | Male | 93 | 0 | 12 | 30 | 1,229.2† | USC | Yes | 3.6 | Yes |
| Prefrontal cortex | ε3/ε3 | Male | 93 | 0 | 3.75 | — | — | USC | Yes | 6.1 | Yes |
| Prefrontal cortex | ε3/ε3 | Male | 76 | 0 | 11.25 | — | — | USC | Yes | 6 | Yes |
| Prefrontal cortex | ε3/ε3 | Male | 86 | 2 | 4.42 | 25 | 48.4 | UCI | Yes | 5.1 | Yes |
| Prefrontal cortex | ε3/ε3 | Male | 83 | 2 | 3.18 | — | — | UCI | Yes | 1.7 | Yes |
| Prefrontal cortex | ε3/ε3 | Male | 86 | 3 | 2.92 | 20 | 37 | UCI | Yes | 3.8 | Yes |
| Prefrontal cortex | ε3/ε3 | Male | 80 | 3 | 4.05 | 26 | 95.1 | UCI | Yes | 4.7 | Yes |
| Prefrontal cortex | ε3/ε4 | Male | 92 | 0 | 5.25 | — | — | USC | Yes | 2.7 | Yes |
| Prefrontal cortex | ε3/ε4 | Male | 90 | 3 | 5.8 | — | — | UCI | Yes | 6.7 | Yes |
| Prefrontal cortex | ε3/ε4 | Male | 97 | 4 | 5.83 | 26 | 7 | UCI | Yes | 5.6 | Yes |
| Prefrontal cortex | ε3/ε4 | Male | 87 | 4 | 6.28 | 22 | 101.5 | UCI | Yes | 4.6 | Yes |
| Cerebellum | ε3/ε3 | Female | 99 | 0 | 9 | 28 | 23.4 | USC | Yes | 8.1 | Yes |
| Cerebellum | ε3/ε3 | Female | 91 | 0 | 8.75 | 20 | 84 | USC | Yes | 8 | Yes |
| Cerebellum | ε3/ε3 | Female | 85 | 0 | 7 | — | — | USC | Yes | 7 | Yes |
| Cerebellum | ε3/ε3 | Female | 95 | 1 | 3.25 | — | — | USC | Yes | 6.1 | Yes |
| Cerebellum | ε3/ε3 | Female | 87 | 2 | 10.75 | 29 | 1 | USC | Yes | 3.8 | Yes |
| Cerebellum | ε3/ε3 | Female | 95 | 2 | 2.92 | — | — | UCI | Yes | 7.9 | Yes |
| Cerebellum | ε3/ε3 | Female | 86 | 3 | 6.17 | 30 | 35.5 | UCI | Yes | 4.9 | Yes |
| Cerebellum | ε3/ε3 | Female | 90 | 3 | 2.2 | — | — | UCI | Yes | 6.7 | Yes |
| Cerebellum | ε3/ε4 | Female | 70 | 1 | 4.17 | 28 | 20.1 | UW | Yes | 1.5 | No |
| Cerebellum | ε3/ε4 | Female | 86 | 3 | 9.33 | 29 | 12.3 | UW | Yes | 4 | No |
| Cerebellum | ε3/ε4 | Female | 97 | 3 | 0 | 26 | 6.6 | UW | Yes | 6.5 | No |
| Cerebellum | ε3/ε4 | Female | 91 | 3 | 11.2 | 23 | 10.1 | UW | Yes | 1.5 | Yes |
| Cerebellum | ε3/ε4 | Female | 85 | 3 | 4.2 | 29 | 37.9 | UW | Yes | 9.3 | No |
| Cerebellum | ε3/ε4 | Female | 89 | 3 | 3.5 | 27 | 28.7 | UW | Yes | 8.1 | No |
| Cerebellum | ε3/ε4 | Female | 74 | 3 | 4.5 | 30 | 44.3 | UW | Yes | 5.1 | No |
| Cerebellum | ε3/ε4 | Female | 91 | 3 | 3.33 | 27 | 42.5 | UCI | Yes | 4.5 | Yes |
| Cerebellum | ε3/ε4 | Female | 87 | 3 | 4.57 | 29 | 85.6 | UCI | Yes | 5.2 | Yes |
| Cerebellum | ε3/ε4 | Female | 93 | 5 | 4.15 | 30 | 81.6 | UCI | Yes | 7.5 | Yes |
| Cerebellum | ε3/ε3 | Male | 82 | 0 | 9 | 28 | — | USC | Yes | 8.6 | Yes |
| Cerebellum | ε3/ε3 | Male | 93 | 0 | 12 | 30 | 1,229.2† | USC | Yes | 2.8 | Yes |
| Cerebellum | ε3/ε3 | Male | 93 | 0 | 3.75 | — | — | USC | Yes | 6.6 | Yes |
| Cerebellum | ε3/ε3 | Male | 76 | 0 | 11.25 | — | — | USC | Yes | 6.8 | Yes |
| Cerebellum | ε3/ε3 | Male | 86 | 2 | 4.42 | 25 | 48.4 | UCI | Yes | 7.5 | Yes |
| Cerebellum | ε3/ε3 | Male | 83 | 2 | 3.18 | — | — | UCI | Yes | 2.9 | Yes |
| Cerebellum | ε3/ε3 | Male | 86 | 3 | 2.92 | 20 | 37 | UCI | Yes | 6.6 | Yes |
| Cerebellum | ε3/ε3 | Male | 80 | 3 | 4.05 | 26 | 95.1 | UCI | Yes | 6.6 | Yes |
| Cerebellum | ε3/ε4 | Male | 92 | 0 | 5.25 | — | — | USC | Yes | 4.1 | Yes |
| Cerebellum | ε3/ε4 | Male | 87 | 1 | 4 | 25 | 8.9 | UW | Yes | 4.4 | No |
| Cerebellum | ε3/ε4 | Male | 90 | 3 | 5.8 | — | — | UCI | Yes | 9.5 | Yes |
| Cerebellum | ε3/ε4 | Male | 97 | 4 | 5.83 | 26 | 7 | UCI | Yes | 7.1 | Yes |
| Cerebellum | ε3/ε4 | Male | 87 | 4 | 6.28 | 22 | 101.5 | UCI | Yes | 8.2 | Yes |

**Supplementary Table 2.**
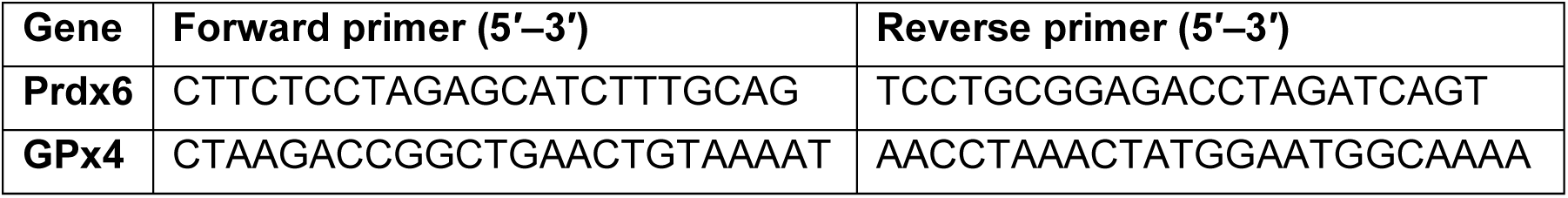
Primer sequences used for Nrf2 ChIP-qPCR. Forward and reverse primers used to amplify EpRE-containing promoter regions of *Prdx6* and *GPx4*.

## Acknowledgements

The authors are grateful to Carol Church and Kymry Jones for assistance with postmortem tissues. The authors appreciate the helpful discussions of brain samples with Carol Miller (USC) and C. Dirk Keene (UW). Tissue for this study was obtained from the USC ADRC Neuropathology Core, National Institute on Aging (NIA)-AG066530. The UCI-ADRC is funded by National Institutes of Health (NIH)/NIA Grant P30-AG066519. E. Head was supported by P30-AG066519. Lab studies were supported by NIH grants to CEF (R01-AG051521, P50-AG05142, P01-AG055367) and Cure Alzheimer’s Fund; R01AG079806 from the NIA, Larry L. Hillblom Foundation Grant 2022-A-010-SUP, the Glenn Foundation for Medical Research and AFAR Grant for Junior Faculty Award, and Navigage Foundation Award to RHS; and T32-AG000037 (PI: Eileen Crimmins) to MAT. Brain specimens were obtained from ADRC Tissue Cores: USC (P50-AG005142, AG066530); University of California Irvine (P30-AG066519); UW (P30-AG066509; U01-AG006781).

